# Mitochondrial Dynamics and Bioenergetics in iPSC-Derived Neurons with Familial Alzheimer’s Disease Mutations

**DOI:** 10.1101/2024.06.24.600414

**Authors:** Courtney MacMullen, Neelam Sharma, Ronald L. Davis

**Author notes:** Lead Contact and Corresponding Author: Ronald L. Davis.

## Abstract

Mitochondrial (MT) dysfunction is a hallmark of Alzheimer’s Disease (AD), but the specific defects across forms of AD are unknown. We measured multiple parameters of MT dynamics and function, and neurite degeneration, in iPSC-derived human neurons possessing natural and engineered mutations in PS1, PS2, and APP genes. Mutations in all three genes altered MT function measured by basal, ATP-linked, and maximal oxygen consumption rate; and spare respiratory capacity, with PS1/PS2 alleles being more severe than APP mutations. Electron flow through Complexes I-IV was decreased in PS1/PS2 mutations but; in contrast, APP alleles had only modest impairments of CI and CII. We measured aspects of MT dynamics including fragmentation, and neurite degeneration, both of which were dramatic in PS1/PS2 alleles, but essentially absent in APP alleles. The marked differences in MT pathology may occur from the distinct ways APP is processed into Aβ and may correlate with the disease severity.

## Introduction

Alzheimer’s Disease (AD) is made up from sporadic forms (sAD) and three major familial forms (fAD). Most AD cases within this spectrum are sporadic^1^, with about 5% of the cases due to mutations in one of three genes: presenilin 1 (PS1), presenilin 2 (PS2), and amyloid beta precursor protein (APP). PS1 and PS2 proteins are the catalytic subunits of γ-secretases, which process APP into a series of amyloid beta (Aβ) peptide fragments. Although the processing of APP produces a diverse set of Aβ peptides, from 37-49 amino acids in length^2^; studies of more than 200 fAD subjects identified a common molecular signature of an increased Aβ42/Aβ40 ratio, with increased Aβ42 abundance as harmful^3–5^. Even though all three genes function in the same pathway of processing APP, some studies have indicated heterogeneity in clinical phenotypes across the three forms of fAD. For instance, a London study evaluated 213 patients with either PS1 or APP mutations and found that PS1 patients had a significantly younger age of onset and a higher incidence of atypical cognitive symptoms apart from memory loss^6^. Despite this clinical heterogeneity, the underlying cellular neuropathology is poorly understood.

The role for mitochondria (MT) in AD and other neurodegenerative diseases has received intense interest in recent years^7, 8^, although the MT cascade hypothesis for AD was proposed two decades ago^9, 10^. These organelles serve in many ways to maintain cell health and have a dynamic life cycle, often referred to by the term “MT dynamics”, which includes MT biogenesis, fission and fusion, MT transport, and removal of damaged MT by mitophagy. The orchestrated processes of MT function and dynamics collectively enable the cell to meet its energy demands, adapt to fluctuations in energy requirements, and maintain overall cellular health. Most importantly, past research using AD post-mortem tissue and cell and animal models of AD have highlighted three broad dysfunctions occurring in neuronal MT: MT fragmentation, and reduced MT content and bioenergetics^11–18^.

However, each of these broad terms reflect extremely complex processes. MT fragmentation is thought to occur from an imbalance between fission and fusion, each involving numerous cellular proteins and a sensing system for monitoring balance. MT content reflects a balance between MT biogenesis and mitophagy, with both processes relying on intricately controlled molecular pathways. MT bioenergetics involves enzymes in the TCA cycle, the electron transport chain (ETC) complexes involved in oxidative phosphorylation, and many other molecules. Thus, our knowledge of MT biology and the MT dysfunction that occurs in AD is too superficial to understand the disease spectrum. For instance, is the broad MT dysfunction identical between different fAD forms and sAD forms, or are there differences that distinguish the forms? In addition, there are many other AD cellular pathologies that are likely triggered by MT dysfunction. Cellular inflammation, synapse failure and loss, axon and dendrite degeneration, increased oxidative stress, excitotoxicity, and other cellular failures may be dependent on the nature of MT dysfunction, producing variation in the types and severity of cellular pathology between individuals with AD^19–21^.

We have begun to address these important questions by systematically detailing the MT dynamics and bioenergetics failures that occur in iPSC-derived neurons representing the 3 major fAD forms. We employed five or seven-iPSC/neuron sets with natural or engineered mutations in PS1, PS2 and APP, and longitudinal assays to follow phenotypic progression. Our results reveal broad MT dysfunction in both MT dynamics and bioenergetics in all three forms, but with marked differences in the specific phenotypes observed and their severity. Remarkably, one broad conclusion is that MT dysfunction is less severe in APP mutants than in PS1 or PS2 mutants, consistent with the lesser clinical severity of APP mutations than PS1 in human subjects.

## Results

### iPSC/neuron sets for the three major forms of fAD

We obtained or generated diseased and control iPSCs as models to study the nature and time course of neuronal MT dysfunction that occurs across the three major forms of fAD and allelic variants within each form. Each of the three forms – PS1, PS2, and APP – is represented by sets of 5-7 iPSC lines used to differentiate glutamatergic neurons (STAR Methods Table 1). Each set contains a naturally occurring allele, derived from a patient with the mutation, along with its isogenic control and for the PS1 and PS2 set, as well as an additional age- and gender-matched control line. Each set also contains the control line Kolf2.1J, characterized as being free of a history for neuropsychiatric disease^22^, and two or three different allelic variants engineered into the line.

The pLVX-UbC-rtTA-Ngn2-2A-AsclI (UNA) plasmid (Figure 1A) was packaged into a lentivirus and used to transduce all lines to create inducible iPSCs that differentiate into neurons upon the addition of doxycycline^23^. Supplemental Figure 1 illustrates one such differentiation; showing that the pluripotency markers, Oct4 and Sox2, are highly expressed in the iPSCs while the neuronal markers of Map2 and Tuj1 were undetectable. By DIV3, the differentiating neurons expressed Map2 and Tuj1 at high levels while Oct4 and Sox2 levels were dramatically decreased. By DIV10, pluripotency marker staining in the neurons was at background levels. At DIV3, the differentiating neurons exhibit short neurite projections that become extensive networks by DIV10 and later.

**Figure 1.**
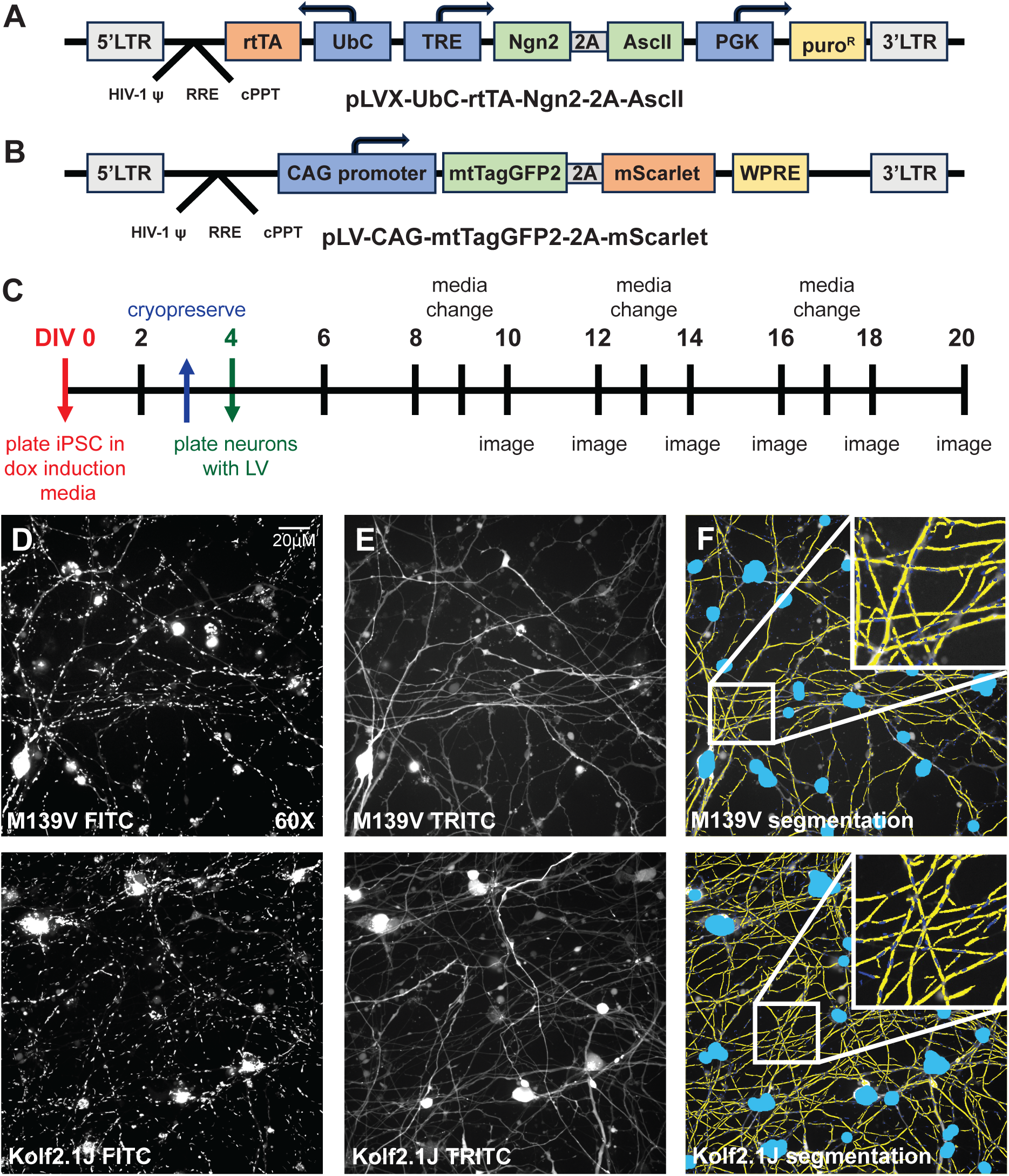
MT dynamics assay overview. **(A)** The pLVX-UbC-rtTA-Ngn2-2A-AscII (UNA) plasmid components that were packaged into a lentiviral vector. **(B)** The pLV-CAG-mtTagGFP2-2A-mScarlet plasmid components that were packaged into a lentiviral vector. **(C)** The MT dynamics assay time course from plating through neuron differentiation, culture, and imaging. Neurons were cryopreserved at DIV3 and replated with lentivirus later at a day designated as DIV4. Images were collected every other day from DIV10-20. **(D)** Representative images of transduced M139V and Kolf2.1J neurons where MT were labeled with TagGFP2. **(E)** Representative images of the same M139V and Kolf2.1J fields where the cytoplasm was labeled with mScarlet. **(F)** Images from (D) and (E) after segmentation of the MT, neurites, and somas. Sky blue areas mark somas and cell debris that were excluded from the analysis. The yellow skeletons represent the segmented neurite projections. The MT are represented as purple dots on top of the yellow neurites and are difficult to see (see inset).

### Multiplexed longitudinal assay for parameters of MT dynamics and neurite complexity

MT are highly dynamic organelles and exhibit multiple actions in living cells that fall under the rubric of MT dynamics, including dividing (fission), fusing (fusion), being newly generated (biogenesis), and degraded (mitophagy). We previously developed phenotypic assays using living primary mouse neurons^24^ and iPSC-derived neurons^25^ that report the balance between fission:fusion and biogenesis:mitophagy, by quantifying average MT length and count, respectively. We modified the CAG>mtTagGFP2 lentivirus originally used for iPSC-derived neurons to label MT to also possess a cytoplasmic reporter, mScarlet, to highlight neurite projections (Figure 1B)^25^. We also modified our original endpoint assays with images collected at a single DIV, to a longitudinal assay with images collected every 2 days between DIV10 and 20 (Figure 1C). DIV10 was the earliest time at which the mtTagGFP2 marker coalesced in the MT so they could be segmented. This modification was made to follow the anticipated progression of MT pathology in the diseased iPSC-derived neurons. Figure 1D-F illustrates representative images of the MT and neurites and their subsequent segmentation for two genotypes under study. The collection of segmented images obtained at each DIV were used to obtain average data for four parameters: soma count, MT length, MT count, and neurite cumulative area (CA). Cumulative area is the average number of pixels representing neurites collapsed across the image fields collected. We normalized the MT count and neurite CA parameters to soma count since there is some neuronal death with time in culture. The parameter MT length was not normalized since this parameter is not correlated with cell density (Supplemental Figure 2).

### MT dynamics pathology in fAD neurons

Figure 2 illustrates the MT dynamics/neurite phenotypes observed in the six PS1 neuron panel plus an age- and gender-matched control for the patient-derived line. We observed significant and progressive MT fragmentation beginning at DIV12, reflected by the MT length parameter, upon comparing data for the AG67 patient-derived AD neurons (PS1 A246E) to the isogenic control ISO-AG67 and the age- and gender-matched control CW10192. No significant difference was observed for this parameter at DIV10. We also observed a significant increase in the MT/soma at DIV16 compared to ISO-AG67 and CW10192, presumably due to MT fragmentation and an imbalance between biogenesis and mitophagy. Finally, we observed significant and progressive neurite degeneration, reflected by neurite CA/soma loss, occurring on DIV20 compared to ISO-AG67 and earlier compared to CW10192.

**Figure 2.**
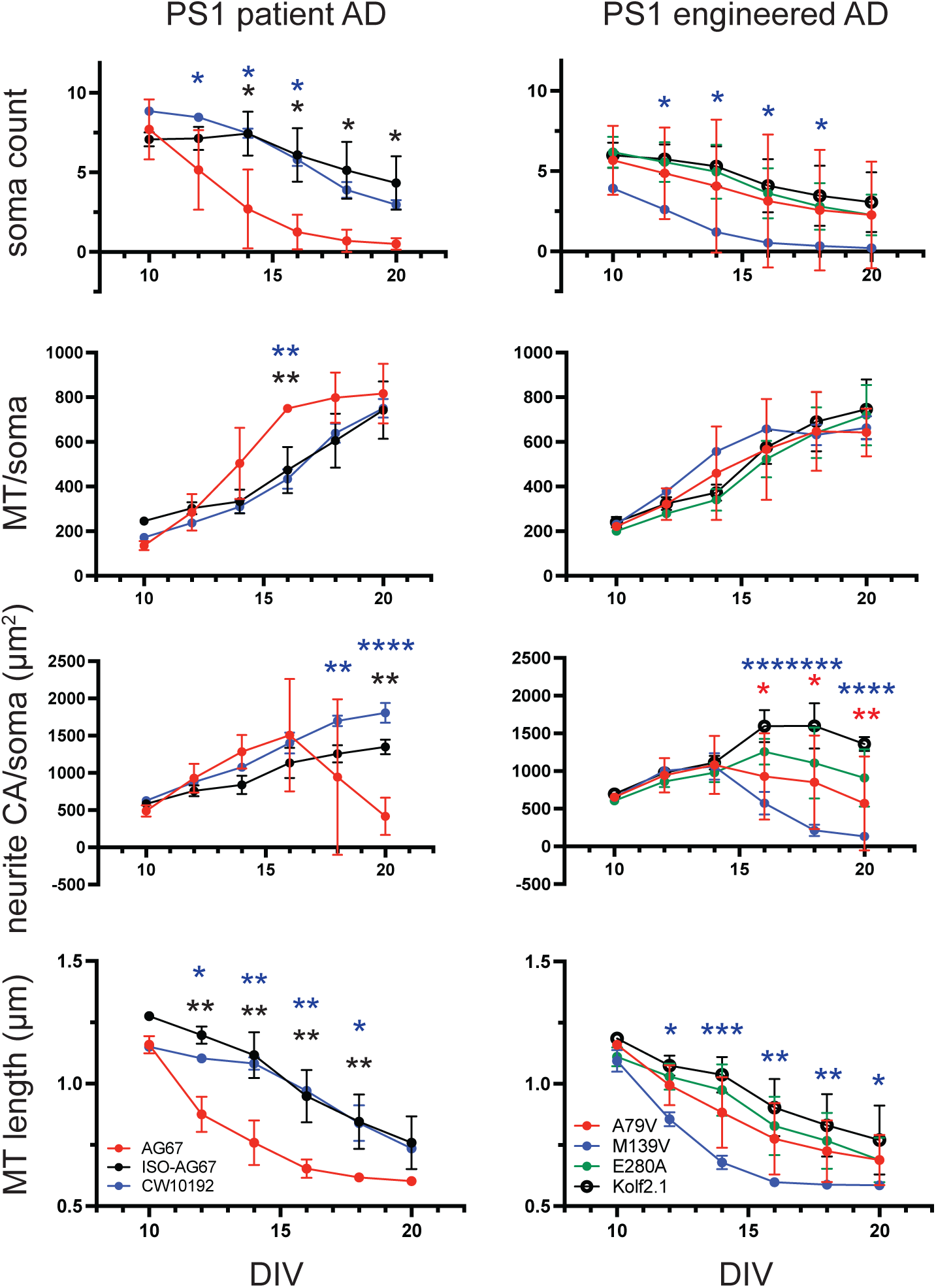
PS1 MT and neurite dynamics. *(Left column)* Soma count, MT/soma, neurite CA/soma and MT length for a PS1 patient iPSC-derived neuron line (AG67:PS1 A246E) along with its isogenic (ISO-AG67) and age- and gender-matched controls (CW10192). The AG67 values (red) were compared to ISO-AG67 (black) and CW10192 (blue). Colored asterisks above each time point indicate significant differences from AG67. The AG67 neurons exhibit soma count loss over time; increased MT/soma at DIV16; decreased neurite CA/soma; and MT fragmentation (length) over the time course relative to ISO-AG67 and CW10192. *(Right column)* Soma count, MT/soma, neurite CA/soma and MT length for three mutant PS1 iPSC-derived neuron lines in which mutations were engineered into the Kolf2.1J control iPSC. The three engineered mutants A79V (red), M139V (blue), and E280A (green) were compared to their isogenic control Kolf2.1J. Colored asterisks above each time point indicate significant differences from Kolf2.1J. M139V had soma count loss, decreased neurite CA/soma, and MT fragmentation. A79V also had a decreased neurite CA/soma ratio over the time course compared to Kolf2.1J *Statistics:* Data represent the average of 3 independent experiments each with 40 technical replicates per genotype. A multifactorial, mixed model ANOVA followed by student t-test *post hoc* analysis was used to analyze the patient AD neurons and its isogenic, and age- and gender-matched control neurons. The same analysis was performed separately on the engineered AD mutant neurons to compare with their isogenic control, Kolf2.1J. *P<0.05, **P<0.01, ***P<0.001, ****P<0.0001.

These data offer several important conclusions that are reinforced, in general, by data presented for the other fAD forms (below). First, the diseased iPSC-derived neurons exhibit MT fragmentation, evidenced by decreased MT length and increased numbers of shortened MT, and neurite degeneration, phenotypes observed in rodent AD models and human postmortem tissue ^12, 26^. Second, the phenotypes observed increase with time in culture as the neurons differentiate. Third, the data suggest that the age- and gender-matched control exhibits significant differences from the isogenic control that may be due to genetic background differences (see Discussion). This observation is reinforced in our analysis of PS2 below.

The recent development of iPSC resources from the iNDI project offered the opportunity to compare allelic variants in PS1, PS2, and APP in a genetic background devoid of known variants for neuropsychiatric disorders. Thus, any phenotypes observed in these neurons can be viewed as being caused solely by the variant at the fAD locus, whereas the phenotypes observed for patient-derived mutants, as with AG67, could be due to the PS1 mutation along with other unknown susceptibility alleles such as APOE status (see STAR Methods Table 1). Among the three alleles assayed, only the PS1 M139V allele exhibited progressive MT fragmentation. PS1 M139V and PS1 A79V showed significantly decreased neurite content from Kolf2.1J at DIVs 16-20. Like AG67, we observed that the fragmentation phenotype occurred earlier than the MT and neurite content phenotypes. Quantitatively, the AG67 and M139V phenotypes appear quite similar, indicating that these may represent complete loss of function alleles.

We also assayed a panel of seven PS2 iPSC-derived neurons and their controls to compare with other fAD forms (Figure 3). The patient derived AG70 (PS2 N141I) neurons exhibited progressive and extreme MT fragmentation beginning at DIV10, reduced MT content per soma after DIV18, and fewer neurites that were detectable after DIV18 (Figure 3). Such phenotypes are not attributable to the advanced age of the donor (81 years of age), since the age- and gender-matched control CW50062 exhibited high neuron survival and increased MT length compared to ISO-AG70 and AG70. The large difference between CW50062 and the isogenic control again emphasizes the importance of making comparisons after controlling for genetic background differences. While AG67 (PS1 A246E) had increased MT/soma at later time points, AG70 displayed decreased MT/soma at DIV18-20. AG70 had very few live somas by DIV18-20; therefore, there were very few MT left to be detected in the assay leading to the decreased MT/soma ratio. Progressive MT fragmentation is the major phenotype observed with the engineered PS2 alleles with two of the three alleles exhibiting this characteristic. Surprisingly, the engineered PS2 N141I allele did not exhibit a significant phenotype in any of the 4 parameters assayed, even though this allele is shared in the patient derived AG70 line. The most parsimonious explanation is that N141I is a mild allele unless in a sensitized genetic background such as AG70. PS2 M239I also had increased MT/soma at later DIVs like AG67 in addition to its fragmentation phenotype. The soma count and neurite CA/soma parameters were negligible or not detected in the engineered alleles.

**Figure 3.**
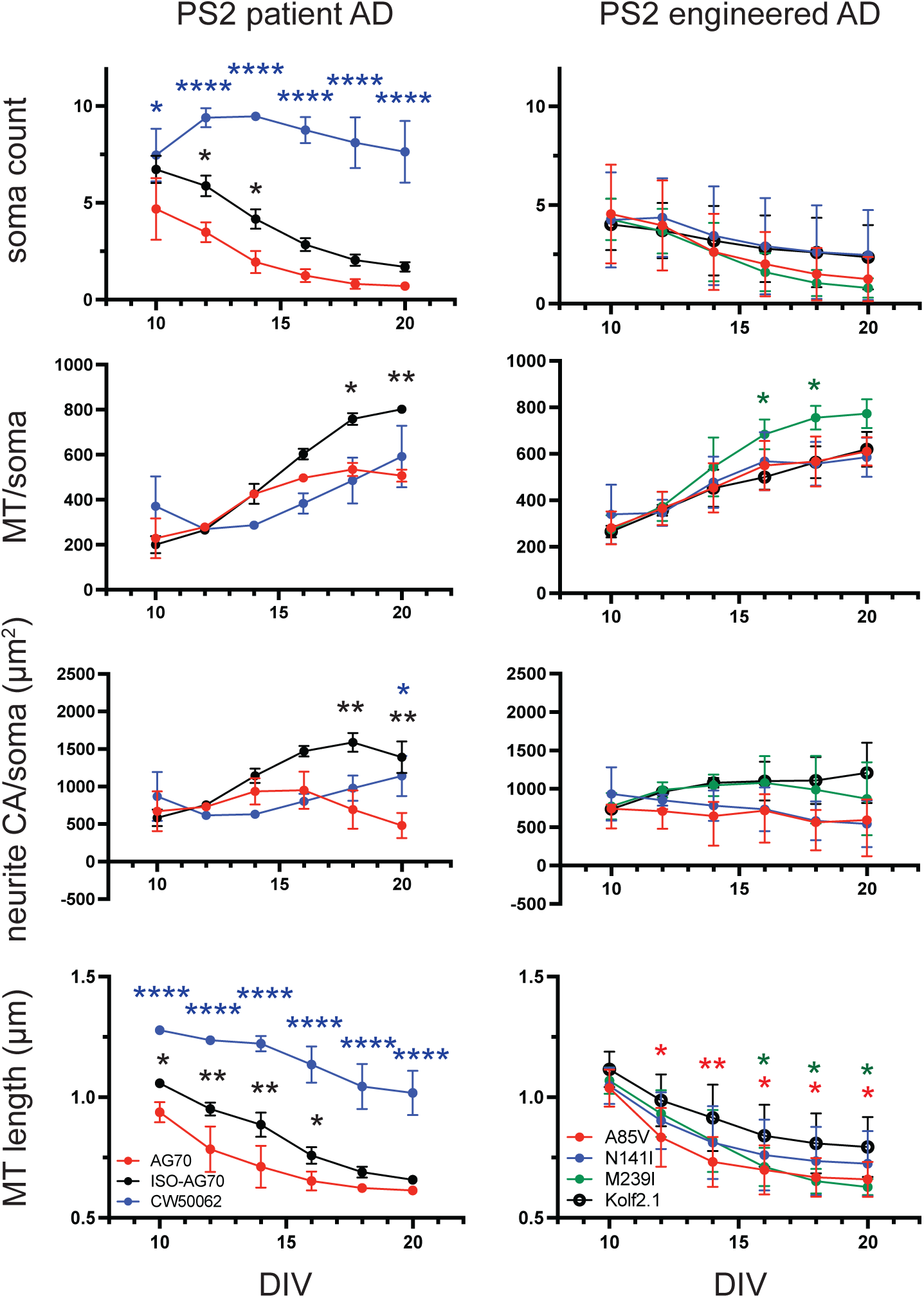
PS2 MT and neurite dynamics. *(Left column)* Soma count, MT/soma, neurite CA/soma and MT length for a PS2 patient iPSC-derived neuron line (AG70:PS2 N141I) along with its isogenic (ISO-AG70) and age- and gender-matched controls (CW50062). The AG70 values (red) were compared to ISO-AG70 (black) and CW50062 (blue). Colored asterisks above each time point indicate significant differences from AG70. The patient AG70 neurons exhibit soma count loss, a decrease in MT/soma, decreased neurite CA/soma and MT fragmentation over the time course relative to ISO-AG70 and CW50062. *(Right column)* Soma count, MT/soma, neurite CA/soma and MT length for three mutant PS2 iPSC-derived neuron lines in which mutations were engineered into the Kolf2.1J control iPSC. The three engineered mutants A85V (red), N141I (blue), and M239I (green) were compared to their isogenic control Kolf2.1J. Colored asterisks above each time point indicate significant differences from Kolf2.1J. The A85V and M239I alleles exhibited significant progressive MT fragmentation. Other phenotypes were modest or undetectable. *Statistics:* Data represent the average of 3 independent experiments each with 40 technical replicates per genotype. A multifactorial, mixed model ANOVA followed by student t-test *post hoc* analysis was used to analyze the patient AD neurons and its isogenic, and age- and gender-matched control neurons. The same analysis was performed separately on the engineered AD mutant neurons to compare with their isogenic control, Kolf2.1J. *P<0.05, **P<0.01, ***P<0.001, ****P<0.0001.

Surprisingly, the panel of five APP iPSC-derived neurons did not show the same MT dynamics phenotypes as the PS1 and PS2 panels. There were no standout differences between KP9 (APP K670N/M671L) neurons and its isogenic control (Figure 4), with only a single difference in MT length at DIV16. This surprise continued with the data obtained from the engineered alleles in that neither the APP E693Q nor the V717I alleles showed any significant differences from Kolf2.1J.

**Figure 4.**
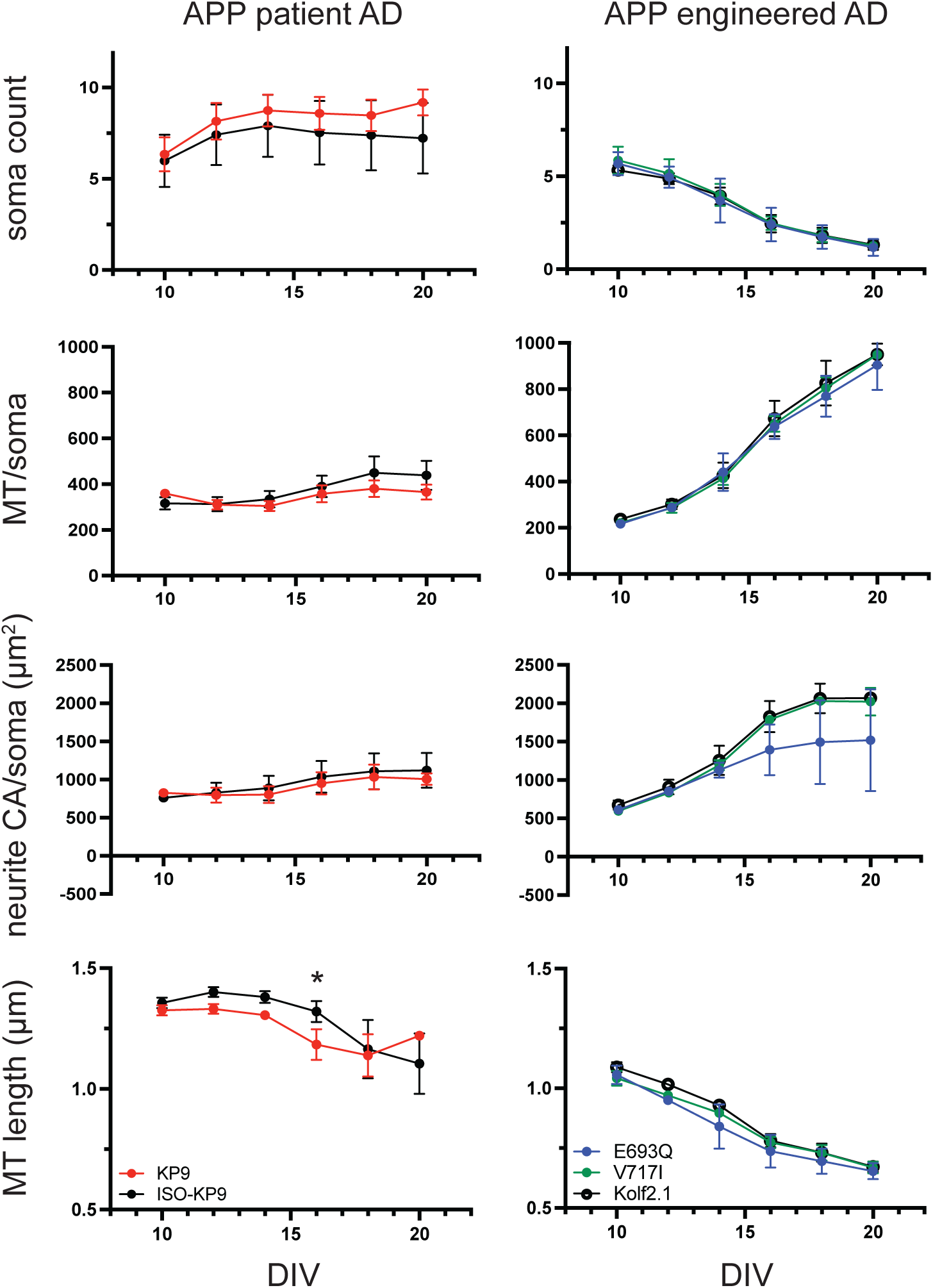
APP MT and neurite dynamics. *(Left column)* Soma count, MT/soma, neurite CA/soma and MT length for an APP patient iPSC-derived neuron line (KP9:K670N/M671L) along with its isogenic control (ISO-KP9). The KP9 values (red) were compared to ISO-KP9 (black). Colored asterisks above each time point indicate significant differences from the control. The patient KP9 neurons exhibit only one timepoint of significant difference for MT length. *(Right column)* Soma count, MT/soma, neurite CA/soma and MT length for two mutant APP iPSC-derived neuron lines in which mutations were engineered into the Kolf2.1J control iPSC. The two engineered mutants E693Q (blue) and V717I (green) were compared to their isogenic control Kolf2.1J. Colored asterisks above each time point indicate significant differences from Kolf2.1J. Neither of the mutant neurons were significantly different from Kolf2.1J for any of the parameters tested. *Statistics:* Data represent the average of 3 independent experiments each with 40 technical replicates per genotype. A multifactorial, mixed model ANOVA followed by student t-test *post hoc* analysis was used to analyze the patient AD neurons and its isogenic, and age- and gender-matched control neurons. The same analysis was performed separately on the engineered AD mutant neurons to compare with their isogenic control, Kolf2.1J. *P<0.05, **P<0.01, ***P<0.001, ****P<0.0001.

We offer more thoughts about this in the Discussion, but MT functional phenotypes were observed in the APP alleles as discussed below.

### MT bioenergetics impairments in familial AD neurons

Mitochondrial stress tests (MST) using the Seahorse analyzer were used to measure and compare possible MT functional phenotypes of PS1, PS2, and APP iPSC-derived neurons. The standard inhibitors of respiration were used at sequential timepoints to measure the neurons’ oxygen consumption rate (OCR). All four measured parameters: basal respiration (Basal), ATP-linked oxygen consumption (ATP), maximal respiration (Maximal), and spare respiratory capacity (SRC) were significantly depressed in the patient-derived AG67 neurons (PS1 A246E) compared to ISO-AG67 at both DIV16 (Figure 5A) and DIV12 (Supplemental Figure 3C; Supplemental Table 1). At DIV8, however, only Basal and SRC were significantly depressed (Supplemental Figure 3A; Supplemental Table 1).

**Figure 5.**
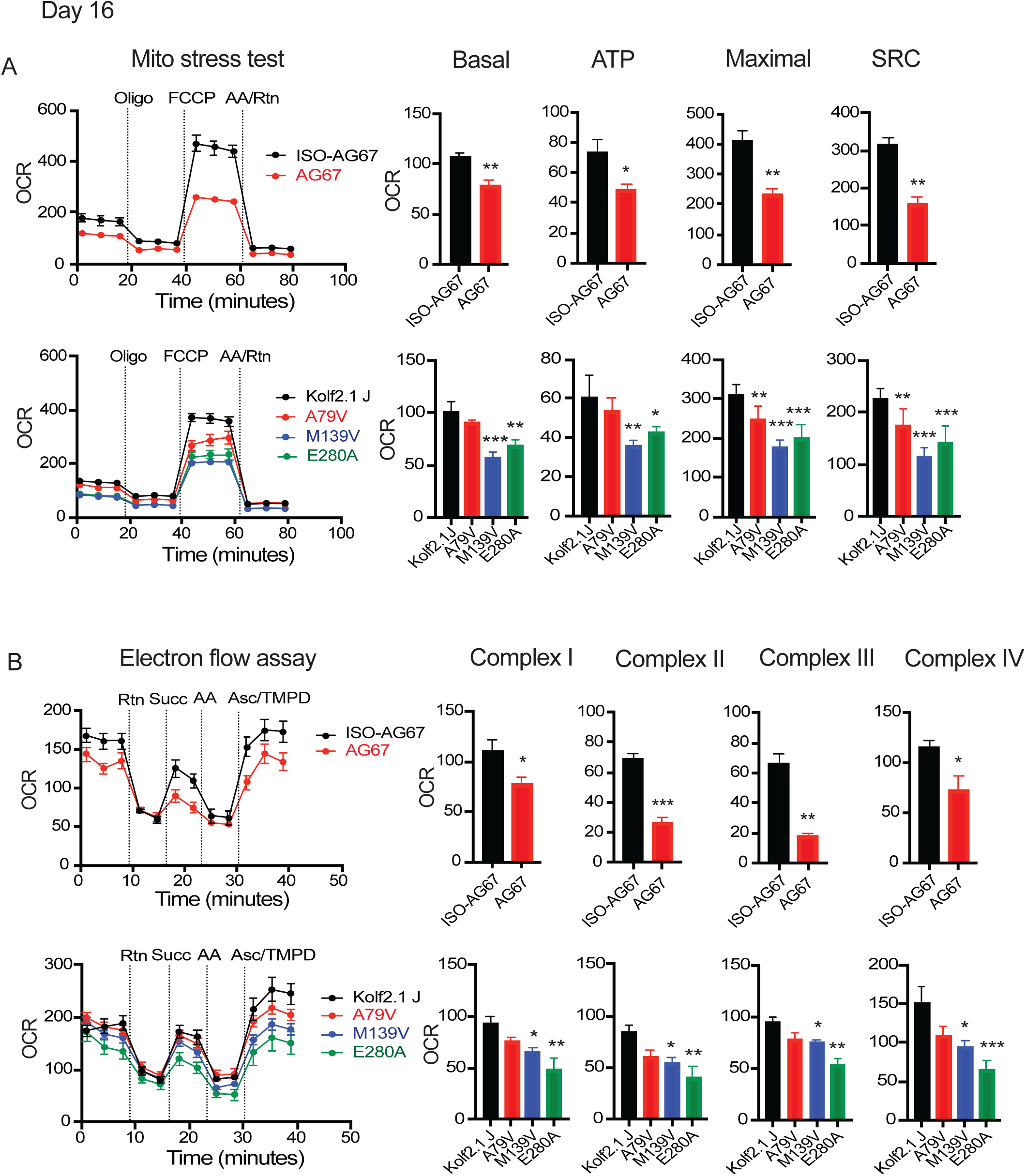
Mitochondrial stress tests and electron flow assays of naturally occurring and engineered PS1 iPSC-derived neurons at DIV16. **(A)** OCR (Oxygen Consumption Rate) was measured in iPSC derived neurons on DIV16 using the Seahorse XF^e^ 96 Analyzer. ETC (Electron Transport Chain) inhibitors were injected into the wells sequentially; Oligomycin=oligo, carbonyl cyanide-p-trifluoromethoxy phenylhydrazone=FCCP, Antimycin A=AA, Rotenone=Rtn. OCR values were normalized to the total number of live neurons/well using calcein staining (pmol/min/live cell number). *Statistics:* Data were analyzed by comparing the average of 12 technical replicates across 3 independent experiments. Statistical significance was calculated using a t-test for ISO-AG67 vs AG67. A one-way ANOVA with Fisher’s *post hoc* test was used for Kolf2.1J vs the 3 engineered alleles. *P<0.05, **P<0.01, ***P<0.001, ****P<0.0001. *(Top row)* Representative line graph of normalized Seahorse data and bar graphs of ISO-AG67 vs AG67 for basal respiration, ATP-linked respiration, maximal respiration, and spare respiratory capacity. *(Bottom row)* Representative line graph of normalized Seahorse data and bar graphs of Kolf2.1J and the 3 engineered alleles for basal respiration, ATP-linked respiration, maximal respiration, and spare respiratory capacity. **(B)** Electron Flow Assay (EFA) measuring the activity of ETC complexes in permeabilized neurons using the Seahorse XF^e^ Analyzer. OCR was measured in permeabilized neurons on DIV16 in response to specific ETC substrates or inhibitors; Rotenone=Rtn, Succinate=Succ, Antimycin A=AA, Ascorbate=Asc, N’-tetramethyl-para-phenylene-diamine=TMPD. The OCR values were normalized to total number of live neurons/well using calcein staining (pmol/min/live cell number). *Statistics:* Data were analyzed by comparing the average of 12 technical replicates across 3 independent experiments. Statistical significance was calculated using a t-test for ISO-AG67 vs AG67. A one-way ANOVA with Fisher’s *post hoc* test was used for Kolf2.1J vs the 3 engineered alleles. *P<0.05, **P<0.01, ***P<0.001, ****P<0.0001. *(Top row)* Representative line graph of normalized EFA OCR data and bar graphs of ISO-AG67 vs AG67 for Complex I, II, III, and IV. *(Bottom row)* Representative line graph of normalized EFA OCR data and bar graphs of Kolf2.1J and the 3 engineered alleles for Complex I, II, III, and IV.

The engineered mutants generally echoed these observations, with at least two of the alleles exhibiting significantly depressed OCR for the four parameters measured at DIV16 (Figure 5A). The A79V was judged to be a mild PS1 allele from the combined data, and M139V was the most severe (Figure 5; Supplemental Figure 3; Supplemental Table 1). Overall, the assays reveal progressively depressed MT function among the PS1 iPSC-derived neurons.

Like PS1, the PS2 patient-derived and engineered mutant neurons also showed MT bioenergetic dysfunction by DIV16 (Figure 6; Supplemental Figure 3; Supplemental Table 1). AG70 patient-derived neurons (PS2 N141I) showed significantly decreased Basal and Maximal values by DIV16, with spare respiratory capacity showing a decreased trend (p=0.067) (Figure 6A). However, these phenotypes were absent at DIV8 and DIV12, except for Maximal at DIV12. The PS2 N141I and PS2 M239I engineered mutants showed decreased OCR values for all 4 parameters at DIV16 (Figure 6A); the A85V allele was less severe with decreased OCR values for Basal and ATP. Some, but not all of the bioenergetic parameters for this set of alleles were decreased at DIV8 and DIV12 (Supplemental Figure 3E, 3G; Supplemental Table 1), indicating some progression in the MT functional pathology with DIV. The relatively severe MT bioenergetics phenotypes for the engineered PS2 N141I neurons is in interesting contrast to the lack of phenotypes in the MT dynamics assay (see Discussion).

**Figure 6.**
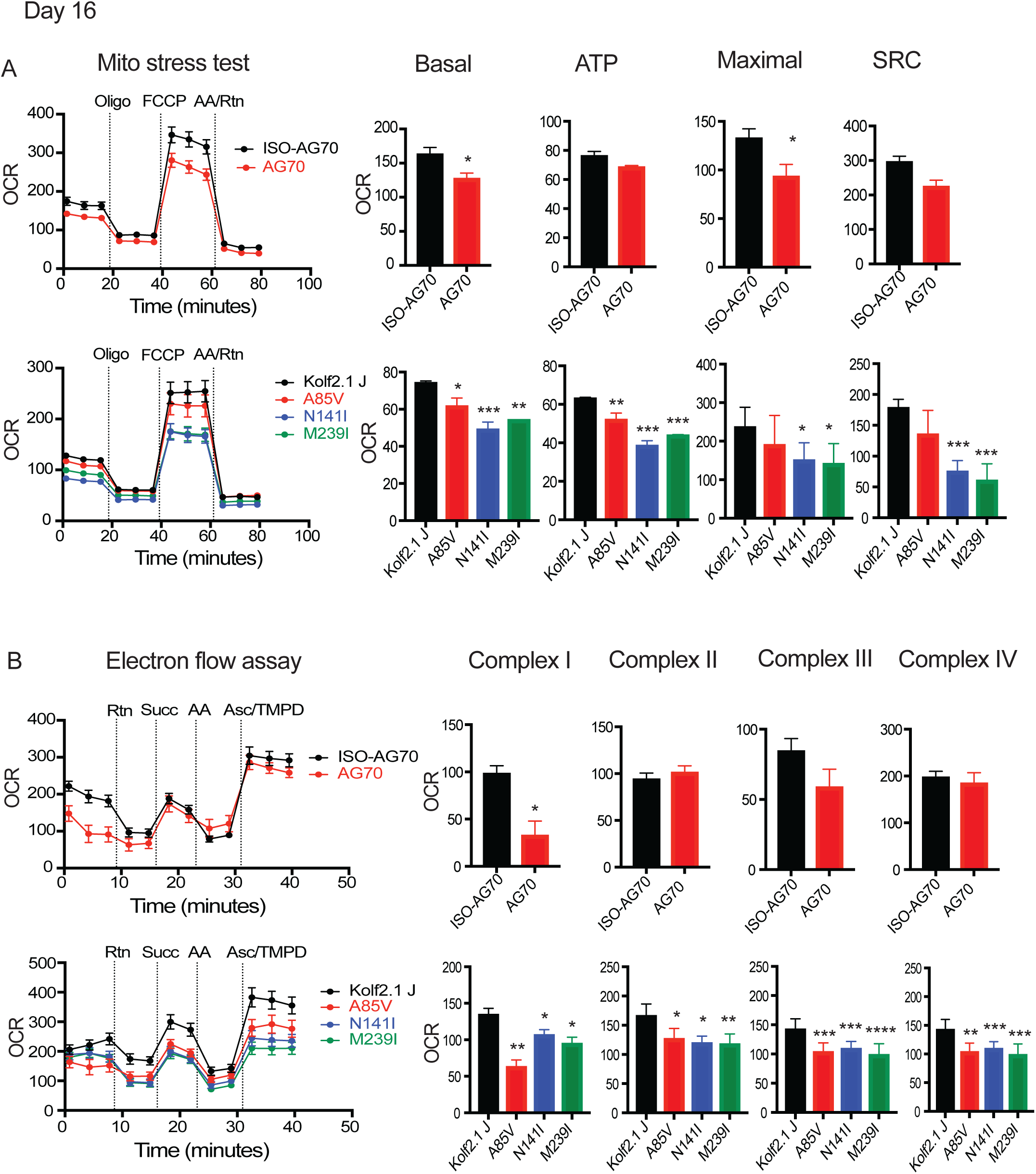
Mitochondrial stress tests and electron flow assays of naturally occurring and engineered PS2 iPSC-derived neurons at DIV16. **(A)** OCR was measured in iPSC derived neurons on DIV16 using the Seahorse XF^e^ 96 Analyzer. ETC inhibitors were injected into the wells sequentially; Oligomycin=oligo, carbonyl cyanide-p-trifluoromethoxy phenylhydrazone=FCCP, Antimycin A=AA, Rotenone=Rtn. OCR values were normalized to the total number of live neurons/well using calcein staining (pmol/min/live cell number). *Statistics:* Data were analyzed by comparing the average of 12 technical replicates across 3 independent experiments. Statistical significance was calculated using a t-test for ISO-AG70 vs AG70. A 2-way ANOVA with Fisher’s *post hoc* test was used for Kolf2.1J vs the 3 engineered alleles. *P<0.05, **P<0.01, ***P<0.001, ****P<0.0001. *(Top row)* Representative line graph of normalized Seahorse data and bar graphs of ISO-AG70 vs AG70 for basal respiration, ATP-linked respiration, maximal respiration, and spare respiratory capacity. *(Bottom row)* Representative line graph of normalized Seahorse data and bar graphs of Kolf2.1J and the 3 engineered alleles for basal respiration, ATP-linked respiration, maximal respiration, and spare respiratory capacity. **(B)** Electron Flow Assay (EFA) measuring the activity of ETC complexes in permeabilized neurons using the Seahorse XF^e^ Analyzer. OCR was measured in permeabilized neurons on DIV16 in response to specific ETC substrates or inhibitors; Rotenone=Rtn, Succinate=Succ, Antimycin A=AA, Ascorbate=Asc, N’-tetramethyl-para-phenylene-diamine=TMPD. The OCR values were normalized to total number of live neurons/well using calcein staining (pmol/min/live cell number). *Statistics:* Data were analyzed by comparing the average of 12 technical replicates across 3 independent experiments. Statistical significance was calculated using a t-test for ISO-AG70 vs AG70. A 2-way ANOVA with Fisher’s *post hoc* test was used for Kolf2.1J vs the 3 engineered alleles. *P<0.05, **P<0.01, ***P<0.001, ****P<0.0001. *(Top row)* Representative experiment line graph of normalized EFA OCR data and three experiment bar graphs of ISO-AG70 vs AG70 for Complex I, II, III, and IV. *(Bottom row)* Representative experiment line graph of normalized EFA OCR data and three experiment bar graphs of Kolf2.1J and the 3 engineered alleles for Complex I, II, III, and IV.

In contrast to the MT dynamics assay, for which the APP mutant neurons showed little or no MT morphological phenotypes, the APP mutant neurons exhibited functional deficits though less severe than PS1 and PS2 mutant neurons. A decreased basal OCR was observed for patient allele KP9 only at DIV16 with no significant differences in other three bioenergetic parameters. The engineered mutants exhibited stronger phenotypes than the patient-derived neurons, with V717I allele exhibiting significantly depressed OCR for all four parameters at DIV16 (Figure 7A). The E693Q allele showed significant decrease only in basal OCR at DIV16. The V717I allele was the most severe, with OCR phenotypes detected as early as DIV8 (Supplemental Figure 3I, 3K; Supplemental Table 1). (see Discussion)

**Figure 7.**
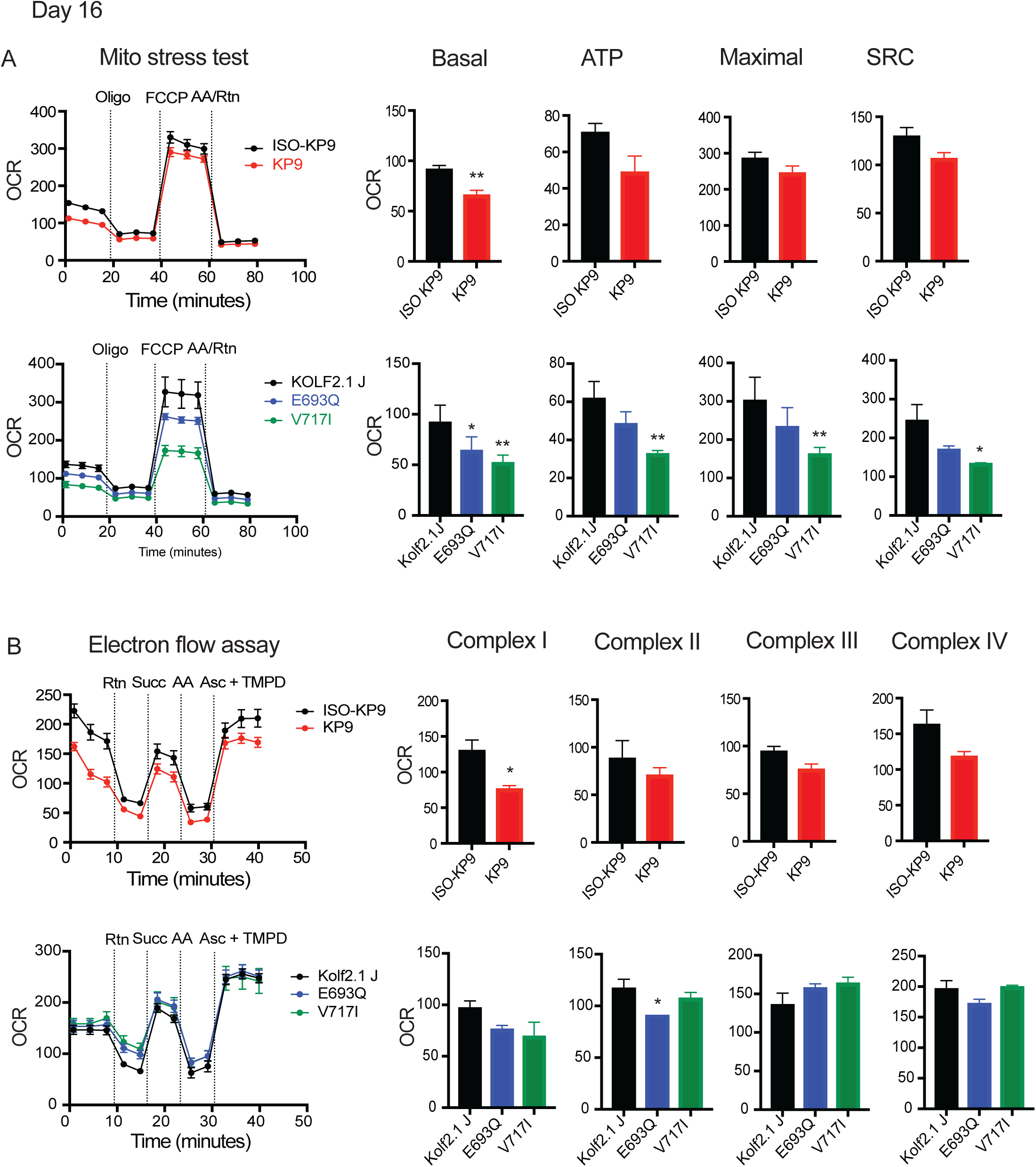
Mitochondrial stress tests and electron flow assays of naturally occurring and engineered APP iPSC-derived neurons at DIV16. **(A)** OCR was measured in iPSC derived neurons on DIV16 using the Seahorse XF^e^ 96 Analyzer. ETC inhibitors were injected into the wells sequentially; Oligomycin=oligo, carbonyl cyanide-p-trifluoromethoxy phenylhydrazone=FCCP, Antimycin A=AA, Rotenone=Rtn. OCR values were normalized to the total number of live neurons/well using calcein staining (pmol/min/live cell number). *Statistics:* Data were analyzed by comparing the average of 12 technical replicates across 3 independent experiments. Statistical significance was calculated using a t-test for ISO-KP9 vs KP9. A 2-way ANOVA with Fisher’s *post hoc* test was used for Kolf2.1J vs the 3 engineered alleles. *P<0.05, **P<0.01, ***P<0.001, ****P<0.0001. *(Top row)* Representative line graph of normalized Seahorse data and bar graphs of ISO-KP9 vs KP9 for basal respiration, ATP-linked respiration, maximal respiration, and spare respiratory capacity. *(Bottom row)* Representative line graph of normalized Seahorse data and bar graphs of Kolf2.1J and the 3 engineered alleles for basal respiration, ATP-linked respiration, maximal respiration, and spare respiratory capacity. **(B)** Electron Flow Assay (EFA) measuring the activity of ETC complexes in permeabilized neurons using the Seahorse XF^e^ Analyzer. OCR was measured in permeabilized neurons on DIV16 in response to specific ETC substrates or inhibitors; Rotenone=Rtn, Succinate=Succ, Antimycin A=AA, Ascorbate=Asc, N’-tetramethyl-para-phenylene-diamine=TMPD. The OCR values were normalized to total number of live neurons/well using calcein staining (pmol/min/live cell number). *Statistics:* Data were analyzed by comparing the average 12 technical replicates across 3 independent experiments. Statistical significance was calculated using a t-test for ISO-AGKP9 vs KP9. A 2-way ANOVA with Fisher’s *post hoc* test was used for Kolf2.1J vs the 3 engineered alleles. *P<0.05, **P<0.01, ***P<0.001, ****P<0.0001. *(Top row)* Representative line graph of normalized EFA OCR data and bar graphs of ISO-KP9 vs KP9 for Complex I, II, III, and IV. *(Bottom row)* Representative line graph of normalized EFA OCR data and bar graphs of Kolf2.1J and the 2 engineered alleles for Complex I, II, III, and IV.

In summary, we observed bioenergetic impairments across all of the three fAD forms and for both patient-derived and engineered mutations. However, there was very significant variability in the strength of the phenotypes among the forms and their allelic variants. If these data translate to the clinical condition, one would predict significant variation in the age-of-onset, progression rate, and/or cognitive processes affected among patients with different familial forms and alleles.

### Differential impairments in electron chain complex activity among familial AD mutant neurons

We also employed the electron flow assay to potentially pinpoint the functional impairments in the electron transport chain (ETC) complexes using permeabilized neurons and the Seahorse analyzer. Complex I and IV, in particular, have been reported to be impacted in AD patients which would affect their ability to meet cellular energy demands ^27, 28^. The activity of each complex was measured by monitoring changes in OCR upon the addition of complex-specific substrates or inhibitors. Initially, pyruvate and malate were present to provide electrons to complex I. Rotenone was then injected to inhibit complex I, preventing electron transfer from NADH to ubiquinone. Succinate was injected next, feeding electrons into the chain via complex II. The third injection was antimycin A, which inhibits complex III, halting electron flow from ubiquinol to cytochrome c. Finally, ascorbate and TMPD were added which donates electrons directly to cytochrome c, bypassing complexes I, II, and III, and allowing the assessment of complex IV activity.

All of the complexes of the patient-derived and the engineered alleles of PS1 had significantly depressed activity at DIV16 except for the A79V allele (Figure 5B), and the phenotypes observed were less severe at earlier assay days (Supplemental Figure 3B, 3D; and Supplemental Table 1). PS2 alleles presented a slightly different picture. Only Complex I was impaired in the patient allele at DIV16 (Figure 6B), but all the engineered alleles were measured as having significantly depressed activity for all of the complexes, like PS1. The phenotypes across the PS2 alleles were less severe at earlier timepoints, but the engineered allele M239I revealed impairments in all complexes at DIV8 except for Complex I (Supplemental Figure 3F, 3H; Supplemental Table 1). The APP alleles showed the mildest effects. Complex I activity was reduced in the patient allele only at DIV16 and some of the engineered alleles exhibited impairments in Complex I and II at DIV12 and 16 (Figure 7; Supplemental Figure 3J, 3L; Supplemental Table 1).

## Discussion

The AD iPSC-derived neurons studied here exhibit the broad MT phenotypes observed in human post mortem tissue and cell and animal models of AD: MT fragmentation and reduced MT content and bioenergetics. The use of human iPSC-derived neurons has the distinct advantages of offering a renewable experimental system, human-derived rather than rodent neurons, and cell populations that can be differentiated into neurons on demand. The overlap of broad MT phenotypes that we observed in the iPSC-derived neurons with other AD model systems offers support for the continued use of iPSC-derived neurons to extract deeper information on MT pathology and other associated neuropathologies in AD.

However, our data, which included measuring MT length, MT count/soma, neurite complexity/soma, cell death via soma count, four parameters of MT oxygen consumption, and electron flow across the four major complexes in 10 alleles of PS1, PS2, and the APP genes, reveal distinct differences in the nature and strength of the phenotypes between the three fAD forms studied. Differences in allelic strength within each gene were expected, because allele strength is influenced by the specific amino acid that is altered and its location in the encoded protein. However, the dramatic differences in phenotypic expression, especially when comparing the engineered alleles in the 3 genes which have the same genetic background, was surprising. For instance, at least some of the engineered PS1 and PS2 alleles show MT fragmentation and neurite degeneration phenotypes, whereas none of the APP alleles displayed these phenotypes across the timecourse (Figures 2-4). DIV16 electron flow assays show that at least some PS1 and PS2 alleles altered electron flow across all four complexes tested, whereas the APP alleles altered only Complex I and Complex II (Figures 5B-7B). All three proteins, APP, PS1, and PS2, are involved in the generation of normal or abnormal protein fragments of APP, with the vast majority of mutations producing the common molecular phenotype of an increased Aβ42/Aβ40 ratio^3, 4^. The direct and relatively simple conclusion from this observation is that AD will ensue with an increased Aβ42/Aβ40 ratio, irrespective of which gene is mutated. However, the proteolytic processing of APP by distinct γ-secretases that use PS1 or PS2 as their catalytic subunit is much more complicated, generating multiple Aβ peptides with lengths of 37, 38, 40, 42, 43, and longer. Thus, the most parsimoneous explanation for the observed phenotypic diversity across the 3 major genes is that different sets of quantitative and qualitative toxic Aβ peptides are produced on a gene and allele specific basis. Consistent with this possibility, Szaruga et al demonstrated that different PS1 mutations impair the enzyme’s sequential proteolytic activity to differing extents, leading to quantitative and potentially qualitative differences in the set of Aβ peptides that are generated^2^. Such differences may be revealed in the clinical presentation of fAD, including the distinct age-of-onset observed between allelic variants^5^. This viewpoint explains the dramatic differences in MT phenotypic expression within and between mutations in the 3 genes.

We also chose to focus on genetic forms of AD for several reasons. Familial forms of AD present relatively early in life, contributing to early-onset cases. We felt that this might translate into more severe MT phenotypes compared to what one might observe with sAD. In addition, the familial forms provided the added certainty that any phenotype detected was due to the mutation. The causes for sAD remain relatively obscure and probably more diverse, resulting from the interactions of biological factors such as the age-associated deterioration of MT interacting with biological factors like genetic susceptibility, chronic inflammation, and environmental factors such as diet, accumulation of toxins, traumatic brain injury, and others. ^29–31^. However, studies in old iNs, directly induced neurons from fibroblasts of donors aged 40 years and older, also illustrated similar MT fragmentation phenotypes as our iPSC-derived AD neurons as well as significantly reduced ETC Complex I transcript expression^32^. Overall, the three panels of AD mutant neurons that form the foundation for our study offered an excellent set to conduct deep investigations into the demise of MT in AD across allelic state for the three major genetic causes of AD.

How do the various types and combinations of Aβ peptides produce MT dysfunction and neurite degeneration? There are two broad possibilities for this that are not mutually exclusive. Multiple studies have reported that APP-derived peptides are transported directly into MT and interact with MT proteins to produce toxic effects^12, 33–38^. The second possibility is that the Aβ peptides sensitize the neuron to the MT dysfunction that occurs with aging, leading to a more precipitous decline in MT function. This possibility views the fAD mutations as susceptibility alleles, similar to other less potent genetic factors that influence AD susceptibility. Other studies have demonstrated that neurite degeneration is a downstream consequence of MT dysfunction. Agents that reduce MT function or content in neurites are known to cause neurite degeneration^12, 39, 40^. The panels of neurons employed may be useful in teasing out the cause and effect relationships between the many molecular and cellular phenotypes observed in AD.

For our analyses of the PS1 and PS2 patient derived alleles, we also employed an age-and-gender matched control with a genetic background distinct from the AD allele and its isogenic control. In the PS1 panel for the MT dynamics assay, the age-and-gender matched control was similar but not identical to the isogenic control (Figure 2). In the PS2 panel, however, the MT dynamics metrics for the age-and-gender matched control were dramatically different from the isogenic control (Figure 3). In addition, the Kolf2.1J isogenic line is a reasonable age-and-gender matched control for the naturally occurring APP mutant (KP9), but its MT metrics are far removed from the KP9 isogenic control (Figure 4). Yet another informative comparison can be made between the naturally occurring PS2 allele N141I and the engineered N141I allele. Only Complex I activity is altered in the naturally occurring allele, whereas Complexes II-IV were measured to be impaired in the engineered allele. These observations emphasize the strong influence of genetic background and the need to use isogenic lines for making the proper comparisons.

The multiplexed, longitudinal assays presented in this study are unique and advantageous in that they allow us to track AD phenotype progression across time in culture. Most studies using AD rodent models and post mortem tissue have employed endpoint assays, which is necessary when using post mortem tissue. However, the data obtained from endpoint assays offer snapshots of AD pathology at restricted timepoints, failing to address pathological progression. Longitudinal assays, using a renewable system such as iPSC-derived neurons, are essential for establishing the linear and/or parallel steps in pathological cascades, such as the mitochondrial cascade hypothesis for Alzheimer’s disease^10^. In addition, the longitudinal assay lends itself nicely to drug discovery efforts, providing an opportunity to administer putative drugs in protective, stalling, or reversal strategies.

## Limitations of study

The study employed relatively pure, iPSC-derived neurons cultured over several weeks to model the neuropathology that occurs in Alzheimer’s disease. This model may or may not reflect all of the complex pathological features that occur in human brain tissue across decades of life.

## Acknowledgements

We wish to thank Michael E. Ward MD, PhD, of the Inherited Neurodegenerative Disease Unit of the NIH/NINDS and William C. Skarnes, PhD, of the Jackson Laboratory for Genomic Medicine, for their assistance in obtaining the Kolf2.1J and engineered mutant iPSC lines used in this study. We would also like to thank Fred H. Gage, PhD and Jerome Mertens, PhD for providing the KP9 line. This study was funded from institutional funds and private philanthropy.

## Author Contributions

Conceptualization; CM, NS, RLD

Methodology: CM, NS, RLD

Data collection: CM, NS

Data analysis and interpretation: CM, NS, RLD

Writing – Original Draft: CM, NS

Writing – Review & Editing: CM, NS, RLD

Supervision: RLD

Funding Acquisition: RLD

## Supplemental Information

Document S1. Supplemental Figures 1-3, Supplemental Figure Legends, and Supplemental Tables 1-3

## STAR Methods

### RESOURCE AVAILABILITY

#### Lead contact

Further information and requests for resources and reagents should be directed to and will be fulfilled by the lead contact, Ronald L. Davis, PhD.

#### Materials availability

Please see STAR Methods Table 1 for cell line origin information. Any cell line that is not commercially available, prohibited from distribution via MTA (iNDI clones), or donated from other investigators (Gage and Mertens) will be available from the lead contact upon request. The pLVX-UbC-rtTA-Ngn2-2A-AsclI (UNA) plasmid is available from Addgene (plasmid# 127288). Any other plasmids and/or reagents not commercially available and generated in this study are available from the lead contact upon request without restrictions.

#### Data and code availability

All data reported in this paper will be shared by the lead contact upon request. Any MetaXpress custom analysis and python codes will be shared by the lead contact upon request. Any additional information required to reanalyze the data reported in this paper is available from the lead contact upon request.

### EXPERIMENTAL MODEL AND STUDY PARTICIPANT DETAILS

#### Cell lines and culture

See STAR Methods Table 1 for information pretaining to each iPSC line used in this study. The AG70, AG67 and KP9 iPSC lines were submitted to Synthego to correct their AD mutations using CRISPR-CAS technology. The corrected iPSC lines were named ISO-AG70, ISO-AG67 and ISO-KP9, respectively. The following iPSC lines were obtained from William C. Skarnes with the iPSC Neurodegenerative Disease Initiative at the Jackson Laboratory for Genomic Medicine (Bar Harbor, ME): Kolf2.1J, PS1 A79V, PS1 M139V, PS1 E280A, PS2 A85V, PS2 N141I, PS2 M239I, APP E693Q and APP V717I ^41^. All engineered mutant Kolf2.1J lines were supplied as early access clones with either heterozygous or homozygous AD mutations. iPSCs were authenticated as to the presence or absence of a specific PS1, PS2, or APP mutation only. Four heterozygous clones were thawed and expanded for each AD mutant. Genomic DNA was extracted from each iPSC clone to confirm the expected genotype. Cell pellets of 2 million iPSC per pellet were isolated and shipped to ThermoFisher Scientific (Waltham, MA) for Scorecard, Karyostat and CellID analysis. A single clone was selected for further analysis from each mutant iPSC line that had correct genotype, appropriate pluripotency marker expression and no large-scale chromosomal abnormalities, see Supplemental Table 2 for the list of early access clones surveyed. iPSCs were transitioned from their original maintenance media and substrate to Essential 8 Flex media and grown on vitronectin coated plates at 37°C in 5% CO2 (see Supplemental Table 3 for original culture conditions). When appropriate, RevitaCell supplement was used in the media at a 1X concentration to prevent apoptosis. iPSCs were clump passaged using 0.5 mM EDTA in 1x Dulbecco’s Phosphate Buffered Saline (DPBS) and single cell passaged using StemPro Accutase. iPSCs were cryopreserved in CTS PSC cryomedium.

**STAR Methods Table 1.**
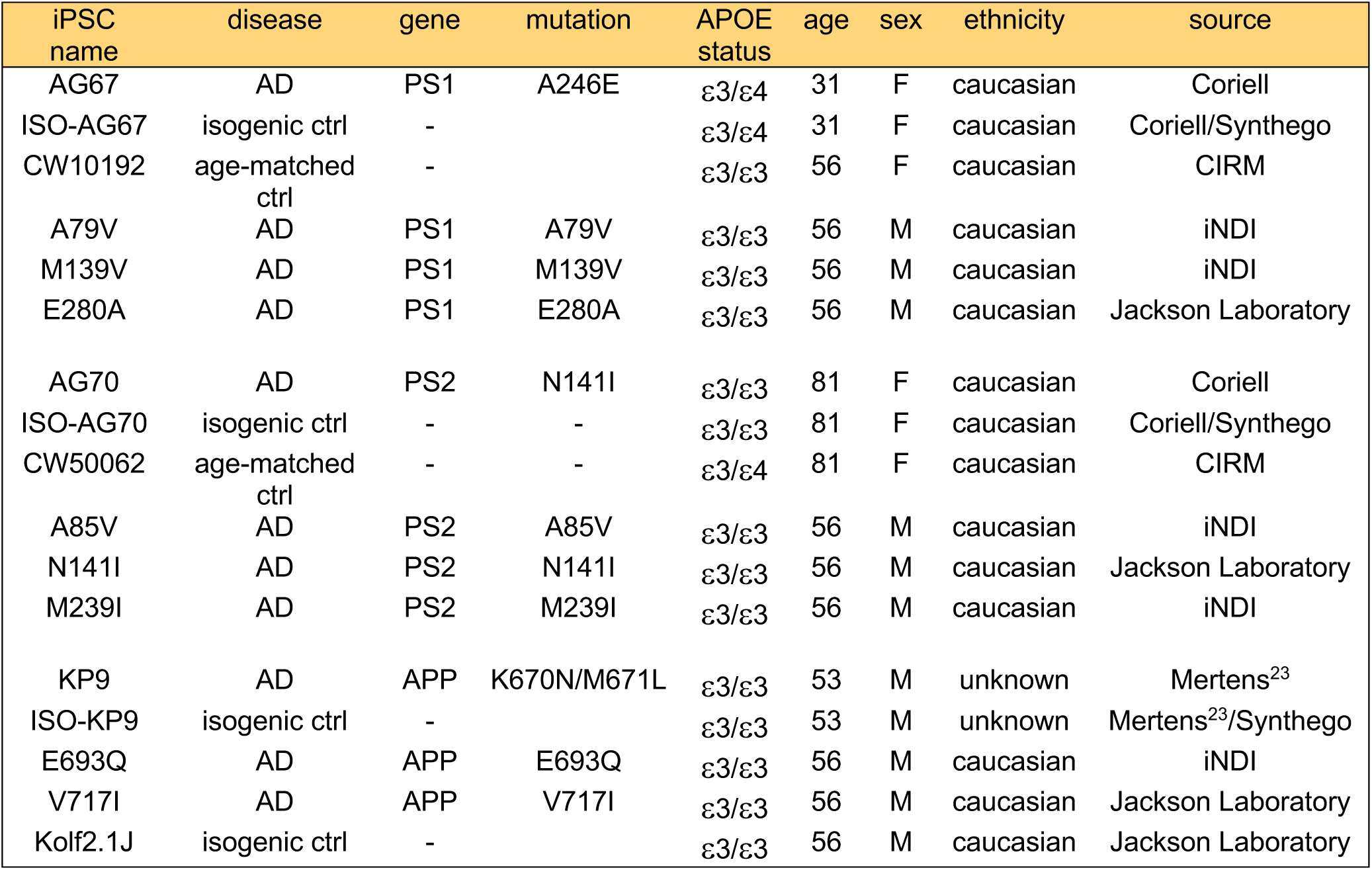

### METHODS DETAILS

#### Transduction of iPSC lines for differentiation

The pLVX-UbC-rtTA-Ngn2-2A-AsclI (UNA) plasmid was packaged into a lentivirus by Vigene Bioscience (Rockville, MD) and used to create inducible iPSC lines^23^. iPSCs were single cell passaged and 175,000 iPSC were transduced with the UNA lentivirus at a multiplicity of infection (MOI) of three in a single well of a 6 well plate. Resulting iPSCs were clump passaged twice and cryopreserved for future use. UNA transduced iPSCs were single cell passaged and 700,000 cells per well in a 6 well plate were selected for transduction positive iPSCs with puromycin at a final concentration of 0.5 μg/mL. Transduction positive iPSCs were maintained in puromycin media for a total of 10 days at which point the iPSCs were cryopreserved for later use.

#### Induction of UNA transduced iPSC

Inducible iPSCs were grown in Essential 8 Flex media on vitronectin coated plates until adequate amounts were available for differentiation. On DIV0, iPSCs were single cell passaged and 6x10^6^ iPSCs were plated in induction media which was comprised of DMEM/F12 base media with final concentrations of the following components: 1X N-2 Supplement, 1X non-essential amino acids (MEM NEAA), and 1x L-glutamine in 15 cm dishes coated with a 1:100 dilution of growth factor reduced matrigel in DMEM/F12 base media. On DIV0 the induction media was supplemented with 1X RevitaCell supplement and 2 μg/mL doxycycline. On DIV1 and DIV2, differentiating cells were washed with 1x DPBS and the induction media was replaced with fresh media containing 2 μg/mL doxycycline. On DIV3, neurons were single cell passaged with StemPro accutase and frozen in 6x10^6^ neuron/cryovial aliquots in 1mL of CTS PSC cryomedium.

#### Pluripotency testing via immunocytochemistry (ICC)

iPSCs were single cell passaged and plated at 12,000 iPSC/well in a 96 well plate coated with a 1:100 dilution of growth factor reduced matrigel in Essential 8 Flex media supplemented with 1X RevitaCell. 100% media changes were performed each day for 2 days with Essential 8 Flex media before beginning the ICC procedure so that colonies could form. Neurons, previously frozen down on DIV3, were thawed and plated on DIV4 in neurobasal plus media which was comprised of neurobasal plus base media with final concentrations of the following components: 1X B-27 plus supplement, 1X GlutaMAX, 1X MEM NEAA, 10 ng/mL brain-derived neurotrophic factor (BDNF), 10 ng/mL NT-3, 1 ug/mL laminin and 1X CultureOne supplement in poly-D-lysine coated black 384w plates at a density of 7,500 neurons/well in a total volume of 80 μl and cultured in a 37°C incubator with 5% CO2. The ICC protocol was performed at the following time points: 3 days after plating for iPSCs and DIV3, DIV10, DIV14 and DIV20 for neurons. Cells were fixed using 4% paraformaldehyde in DPBS for 15min. Cells were washed twice with 50% DPBS reagent changes and permeabilized with 0.5% triton X-100 in DPBS for 15 min. Cells were washed twice with 50% DPBS reagent changes and then blocked in 3% bovine serum albumin (BSA) in DPBS for 1 hr at room temperature. Cells were washed twice with 50% DPBS reagent changes and primary antibodies (rabbit polyclonal Map2, mouse monoclonal beta-III-tubulin, rabbit polyclonal Oct4, and mouse monoclonal Sox2) were added at a 1:500 dilution in BSA blocking buffer and incubated at 4°C overnight. Cells were washed twice with 50% DPBS reagent changes and secondary antibodies (Alexa fluro 488 goat anti mouse and alexa fluro 568 goat anti rabbit) were added at a 1:1000 dilution in BSA blocking buffer and incubated for 1 hr at room temperature without exposure to light. Cells were washed twice with 50% DPBS reagent changes and were then imaged in 2D using the FITC and TRITC channels of a Molecular Devices (San Jose, CA) Image Xpress Micro Confocal High-Content Imaging System. iPSC images were captured using the 20X objective while neuron images were captured using the 60X objective.

#### MT dynamics assay

The pCAG-mtTagGFP2-2A-mScarlet plasmid was packaged into a lentivirus by SignaGen Laboratories (Frederick, MD). Neurons were thawed and plated on DIV4 with LV-CAG-mtTagGFP2-2A-mScarlet at a MOI=3 in neurobasal plus media supplemented with 1X CultureOne in poly-D-lysine (PDL) coated black 384w plates at a density of 15,000 neurons/well in a total volume of 40 μl. A total of 40 replicate wells per neuron type were plated in each experiment. A total of 3 experiments were plated for each neuron set, each experiment beginning on a different day to maintain independence. On DIV5, 20 μl of the media was removed and an additional 60 μl of fresh neurobasal plus media supplemented with 1X CultureOne was added bringing the final volume to 80 μl. Every 4 days, 50% media changes were performed with neurobasal plus media supplemented with 1X CultureOne until DIV13 where CultureOne was no longer added to the media. Beginning at DIV10, each experiment was imaged every two days until DIV20. Images were captured using the Image Xpress Micro Confocal High-Content Imaging System with 60X objective, 0.95 NA with 60μm pinhole to image 4 fields per well. Each field was captured in a 5 step Z-series (step size 0.7μm) and resulting images were collapsed into a 2D maximum intensity projection image for analysis. A custom segmentation algorithm was applied to each image using the MetaXpress software v.6.5.4.532 where MT and neurite extensions were labeled. The following parameters were collected from these images: soma count, total MT count, neurite cumulative area (CA), and median MT length for all MT in image. Custom python codes extracted the MT and neurite data from these field images and applied an IQR (inter quartile range) outlier removal to remove any irregular fields. The MT count and neurite CA parameters were divided by the soma count on a field basis and the 4 fields per well were averaged into average counts per well. The median MT length and soma CT values were not normalized. Experimental average data for each parameter were determined per neuron type from the 40 technical replicates.

#### MT length assay

To illustrate that the MT length parameter was independent of soma count, we plated Kolf2.1J isogenic control neurons in the same manner as described in the MT dynamics assay; however, we varied the density of cells plated. Twenty-four wells of the following cell densities were plated, all with the same pCAG-mtTagGFP2-2A-mScarlet lentivirus MOI=3: 20K, 17.5K, 15K, 12.5K and 10K cells per well. Images were collected and processed from a single time point (DIV16) as described above. The same MT and neurite parameters were collected and analyzed as described above.

#### MT bioenergetics assay

Seahorse plates were coated with 60 μL of 0.1 mg/mL poly-D-lysine (PDL) overnight at 37°C. Plates were washed three times with sterile water and air-dried for 1 hr followed by an additional 1 hr incubation in a 37°C incubator. Neurons, frozen at DIV3, were thawed and plated on PDL coated Seahorse plates at a density of 30,000 cells/well in a total volume of 180 μL of neurobasal plus medium supplemented with 1X CultureOne. A total of 12 replicate wells per neuron type were plated in each experiment. 50% media changes were performed every 4 days until the assay was performed. The mito-stress tests (MST) or electron flow assays (EFA) were performed on DIV8, DIV12, and DIV16 using a Seahorse XF^e^96 analyzer. All experiments were repeated three times with neurons plated on different days.

#### Mito-stress test

Neurons were washed 1X with assay media comprised of XF base medium, 10 mM glucose, 2 mM glutamine, and 1 mM pyruvate. Calcein was then added to each well of the assay plate at a final concentration of 10 µM. The assay plate was incubated at 37°C within a controlled environment, devoid of CO2, for 30min. After this incubation, the plate was imaged using the InCell 6000 analyzer (GE Healthcare Technologies, Chicago, IL) at 37°C to obtain live cell counts for calcein normalization. Live cells were counted using a customized segmentation protocol using the InCell Developer 1.9.2 software. During the 30 min incubation, a hydrated Seahorse cartridge was loaded with specific inhibitors: port A – 2 µM oligomycin, port B – 4 µM carbonyl cyanide p-(trifluoromethoxy) phenylhydrazone (FCCP), and port C – 1 µM rotenone and 1 µM antimycin A. The cartridge was loaded into the analyzer and a calibration was performed. After calcein imaging, the plate was inserted into the XF^e^96 analyzer and an equilibration was performed followed by the comprehensive MT stress test. For each injection, 3 oxygen consumption rate (OCR) measurements were taken. Four mito stress test parameters were assayed: basal respiration (Basal), the general resting state respiratory capacity of the neurons; maximal respiration (Maximal), the amount of energy released when challenged with the electron transport chain uncoupler FCCP; spare respiratory capacity (SRC), the difference between maximal and basal respiration values which indicates the neuron’s energy reserve under periods of stress; and ATP-linked OCR (ATP), an estimation of the amount of energy able to be produced by neurons which is measured after the oligomycin injection. Average OCR well values were normalized to the total number of live cells in each well using the Wave software v2.6.3; this normalized data was then exported into the XF test report generator of the Wave software. Average values of the 12 replicate wells were derived for each experiment for the following MT respiratory parameters: Basal, Maximal, SPR and ATP.

#### Electron flow assay

90 µL of neurobasal plus media was removed from each assay well and replaced with fresh media supplemented with 10 µM final concentration calcein followed by a 30 min incubation at 37°C. Plates were imaged to obtain live cell counts for normalization as described above. During imaging, a hydrated Seahorse cartridge was loaded with substrates and inhibitors tailored for specific cellular responses: port A – 2 µM rotenone, port B – 10 mM succinate, port C – 2 µM antimycin A, and port D – 10 mM ascorbate and 100µM N,N,N′,N′-tetramethyl-para-phenylene-diamine (TMPD), followed by a calibration step. After imaging, neurons were permeablized by washing once with 1X MAS buffer, comprised of 70 mM sucrose, 220 mM mannitol, 10 mM potassium phosphate monobasic (KH2PO4), 5 mM magnesium chloride hexahydrate (MgCl2), 2 mM 4-(2-hydroxyethyl)-1-piperazineethanesulfonic acid (HEPES), and 1 mM ethylene glycol tetraacetic acid (EGTA). Then 1X MAS assay buffer supplemented with 0.2% BSA, 10 mM pyruvate, 1 mM malate, 4 mM Adenosine diphosphate (ADP), and 1 nM XF plasma membrane permeabilizer (PMP) reagent was added to the neurons, the plate was loaded into the XF^e^96 analyzer, and the assay was run. Two measurements were recorded for each injection which were then averaged per well and normalized to the total number of live cells in each well as described above. Average values of the 12 replicate wells were derived for each experiment to obtain individual ETC complex activity levels.

### QUANTIFICATION AND STATISTICAL ANALYSIS

#### MT dynamics assay

The averaged data for each MT and neurite parameter assessed from each of the three experiments were then averaged together for each time point assayed. A multifactorial, mixed model ANOVA was performed using the standard least squares model in JMP v15, with experimental replication, day, neuron, and neuron crossed with day, being fixed factors. Subject, a combination of neuron with specific replication, was assigned a random effect. *Post hoc* analysis was performed with Student t-test. For each gene, patient-derived AD neurons and their age- and gender-matched and isogenic controls were included in a single ANOVA while the engineered AD mutants and their isogenic control, Kolf2.1J, were included in a separate ANOVA.

#### MT length assay

A single experiment was performed for this assay. The averaged well data from the 24 replicate wells for each cell density was compared for each MT and neurite parameter. One way ANOVA with *post-hoc* Fisher’s test was performed to derive statistical significance using Prism v10.1.1.

#### Mito-stress test

Average values of the 12 replicate wells were derived for each of the three independent experiments for the following MT respiratory parameters: Basal, ATP, Maximal, and SRC. One way ANOVA with *post-hoc* Fisher’s test or unpaired t-test was performed to derive statistical significance using Prism v10.1.1. Patient-derived AD neurons and their isogenic controls were included in the ANOVA while the engineered AD mutants and their isogenic control, Kolf2.1J, were included in a separate ANOVA.

#### Electron flow assay

Average values of the 12 replicate wells were derived for each experiment to obtain individual ETC complex activity levels. One way ANOVA with *post-hoc* Fisher’s test or unpaired t-test was performed to derive statistical significance using Prism v10.1.1 (Graphpad, Boston, MA). Patient-derived AD neurons and their isogenic controls were included in the ANOVA while the engineered AD mutants and their isogenic control, Kolf2.1J, were included in a separate ANOVA.

