## Supplemental Figures and Tables for "Mitochondrial Dynamics and Bioenergetics in iPSC-Derived Neurons with Familial Alzheimer’s Disease Mutations"

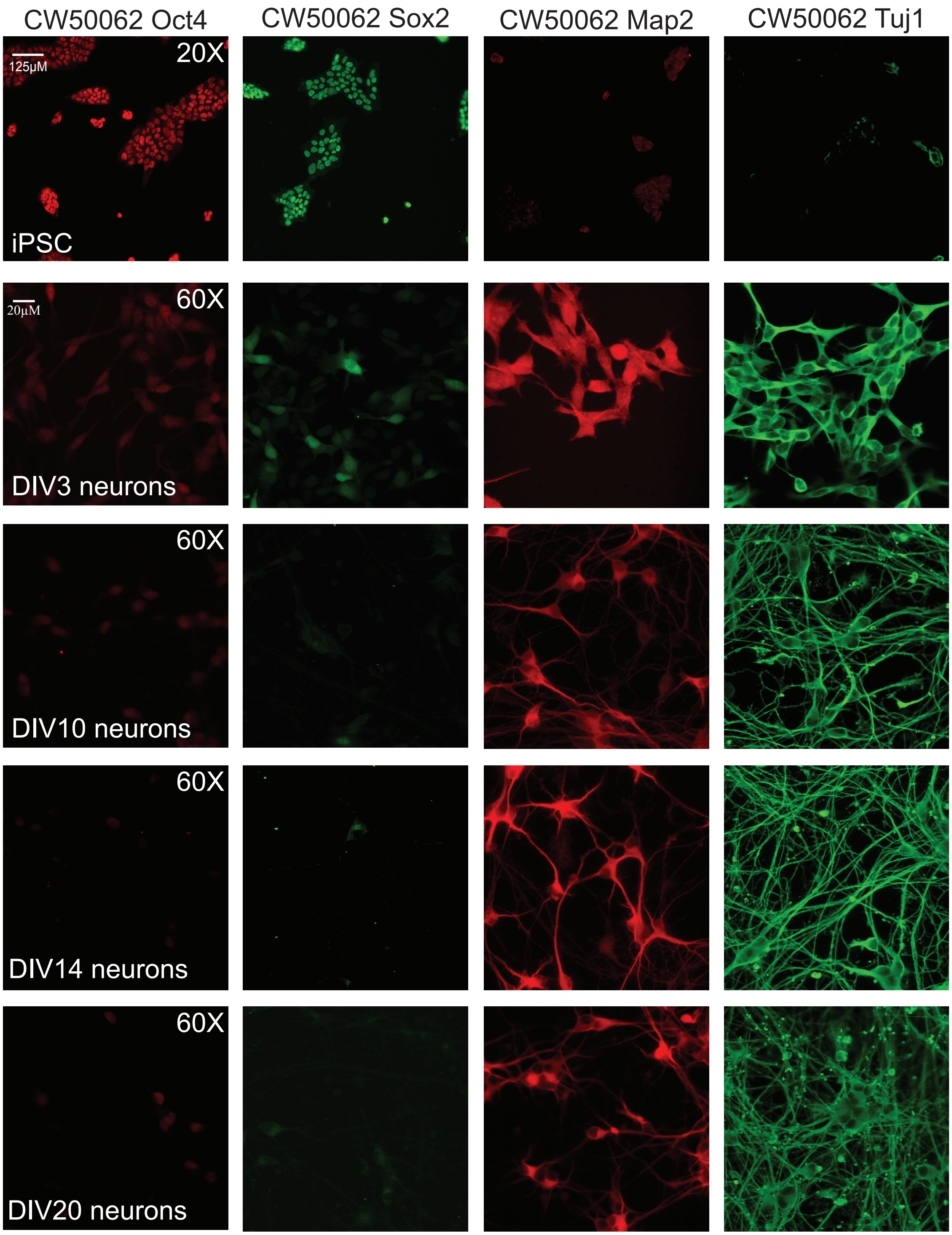

Supplemental Figure 1

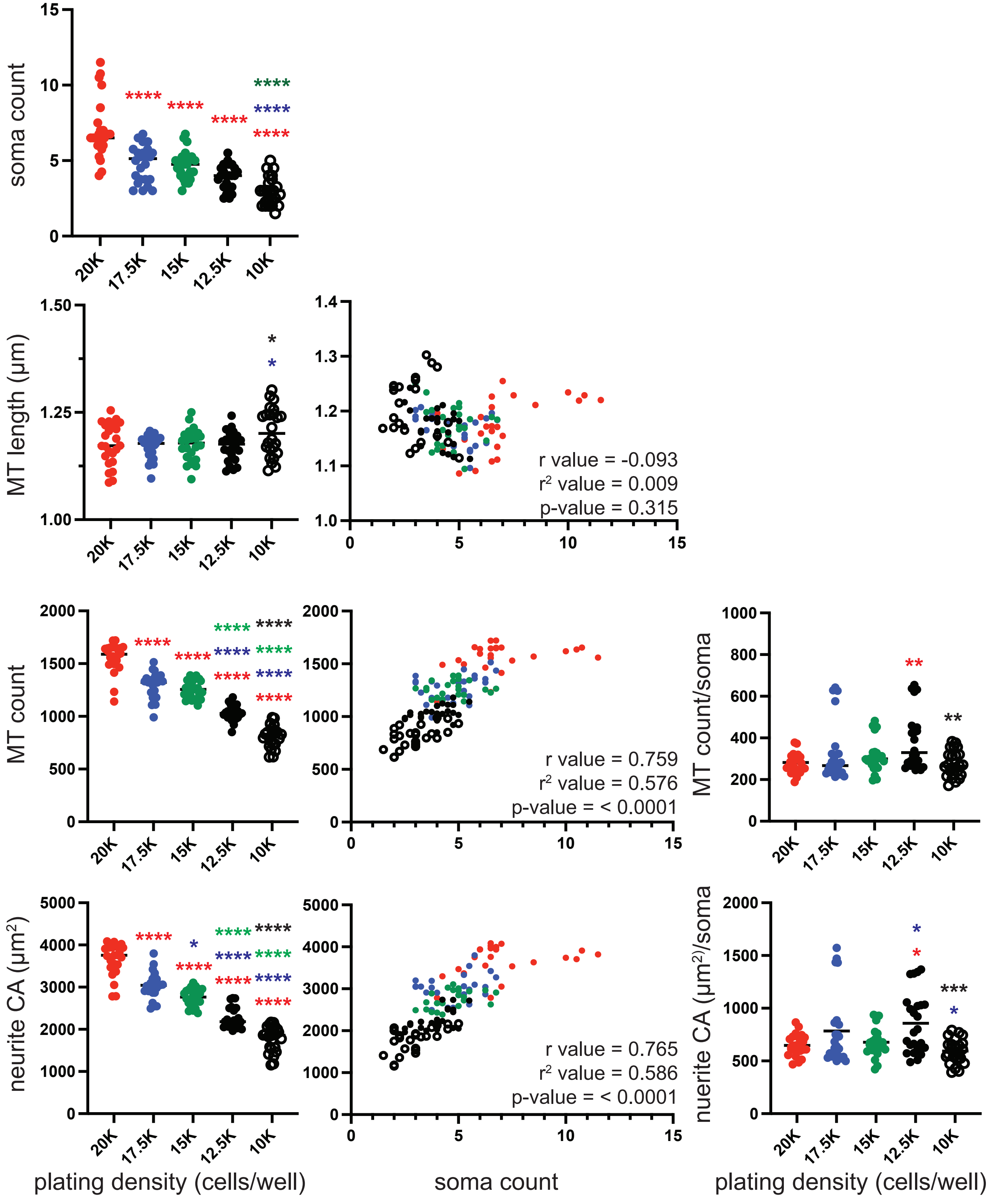

Supplemental Figure 2

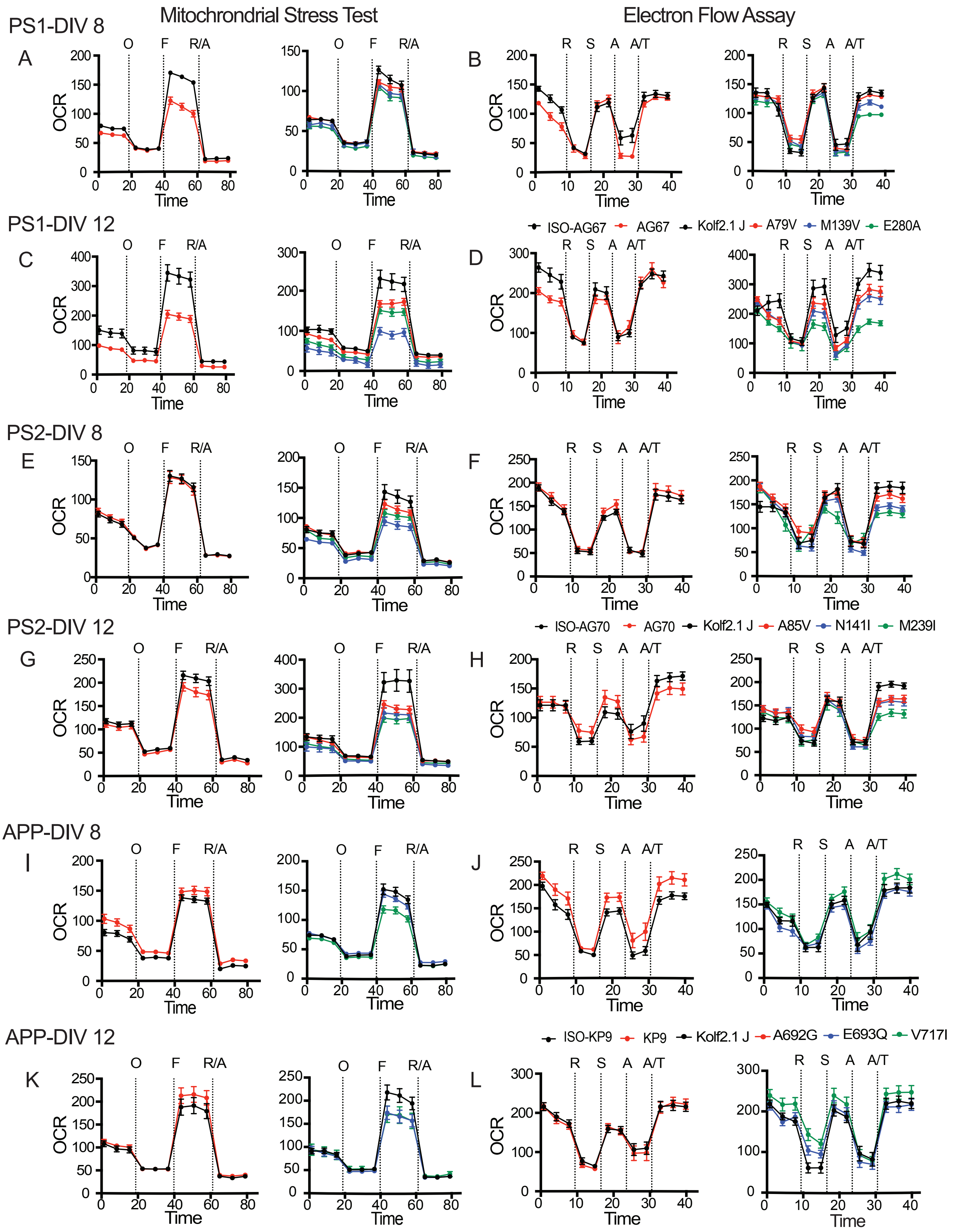

Supplemental Figure 3

### Supplemental Figure Legends

**Supplemental Figure 1. Antigenic marker expression in iPSC and differentiated neurons.**
The pluripotency markers of Oct4 and Sox2 were expressed at high levels in the CW50062 iPSCs. Neuronal differentiation was followed at DIV3, DIV10, DIV14, and DIV20. During differentiation, Oct4 and Sox2 pluripotency marker expression decreased significantly and the neuronal markers Map2 and Tuj1 became highly expressed. DIV3 neurons have very short neurite projections compared to later time points as evidenced by the Tuj1 staining. The very low level of signal in the iPSCs using neuronal antigenic markers and in the differentiated neurons using pluripotency antigenic markers is at background levels.

**Supplemental Figure 2. MT length is independent of soma count**
Isogenic control Kolf2.1J neurons were plated at the following seeding densities per 384 well (20K in red, 17.5K in blue, 15K in green, 12.5K in filled black, and 10K in open black circles) and were imaged on DIV16. Soma count, MT length, MT count, and neurite CA parameters were graphed for each cell density. All parameters except for MT length were dependent on the number of cells plated per well. Correlation graphs comparing MT length to soma count, MT count to soma count, and neurite CA to soma count were also generated where each of the 24 wells plated were colored according to their cell density. These graphs illustrate that MT count and neurite CA are positively correlated with soma count while there is no correlation between MT length and soma count. In addition, the data for MT count and neurite CA were normalized to soma count with no difference found in these parameters at the 20K, 17.5K, and 15K cell plating densities. Statistics: We used 24 wells per cell density and analyzed the data with a one-way ANOVA using multiple comparisons for the soma count, MT length, MT count, and neurite CA parameters. A Pearson correlation analysis was performed and r, $r^2$, and resulting p-values are represented on the graphs. MT count/soma and neurite CA/soma values were also analyzed with a one-way ANOVA using multiple comparisons. *P<0.05, **P<0.01, ***P<0.001, ****P<0.0001. Colored asterisks indicate a significant difference between data points representing that cell density and the one at which the asterisks are located.

**Supplemental Figure 3. Mitochondrial stress test and electron flow assays of naturally occurring and engineered AD iPSC-derived neurons.**
(Mitochondrial Stress Test; left columns) OCR was measured in AD iPSC derived neurons on DIV8 and DIV12 using the Seahorse $XF^e$ 96 Analyzer. ETC Inhibitors were injected into the wells sequentially; Oligomycin=O, carbonyl cyanide-p-trifluoromethoxy phenylhydrazone=F, Rotenone=R, Antimycin A=A. OCR values were normalized to the total number of live neurons/well using calcein staining (pmol/min/live cell number). Three independent experiments were averaged together and illustrated in the line graphs. The statistics for these line graphs are found in Supplemental Table 1.
(Electron Flow Assay; right columns) OCR was measured in permeabilized AD iPSC-derived neurons on DIV8 and DIV12 in response to specific ETC substrates or inhibitors; Rotenone=R, Succinate=S, Antimycin A=A, Ascorbate plus N'-tetramethyl-para-phenylene-diamine=A/T. The OCR values were normalized to total number of live neurons/well using calcein staining (pmol/min/live cell number). Three independent experiments were averaged together and illustrated in the line graphs. The statistics for these line graphs are found in Supplemental Table 1.

| | | P-Values for Mitochondrial Stress Test | | | | P-Values for Electron Flow Assay | | | |
|---|---|---|---|---|---|---|---|---|---|
| | **Genotype** | **Basal respiration** | **ATP linked OCR** | **Maximal respiration** | **Spare respiratory capacity** | **Complex I** | **Complex II** | **Complex III** | **Complex IV** |
| Presenilin 1 | **Day 8** | | | | | | | | |
| | ISO AG67 | | | | | | | | |
| | AG67 | ≤0.05 | n.s. | n.s. | ≤0.05 | ≤0.05 | n.s. | n.s. | n.s. |
| | Kolf2.1J | | | | | | | | |
| | A79V | n.s. | n.s. | n.s. | n.s. | n.s. | n.s. | n.s. | n.s. |
| | M139V | ≤0.05 | n.s. | ≤0.05 | ≤0.05 | n.s. | n.s. | n.s. | ≤0.05 |
| | E280A | n.s. | n.s. | n.s. | n.s. | n.s. | n.s. | n.s. | ≤0.01 |
| | **Day 12** | | | | | | | | |
| | ISO AG67 | | | | | | | | |
| | AG67 | ≤0.05 | ≤0.05 | ≤0.05 | ≤0.01 | ≤0.05 | n.s. | n.s. | n.s. |
| | Kolf2.1J | | | | | | | | |
| | A79V | ≤0.01 | ≤0.05 | ≤0.05 | n.s. | n.s. | n.s. | n.s. | n.s. |
| | M139V | ≤0.001 | ≤0.001 | ≤0.001 | ≤0.01 | n.s. | ≤0.05 | ≤0.05 | n.s. |
| | E280A | ≤0.001 | ≤0.001 | ≤0.05 | n.s. | ≤0.01 | ≤0.01 | ≤0.01 | ≤0.001 |
| Presenilin 2 | **Day 8** | | | | | | | | |
| | ISO AG70 | | | | | | | | |
| | AG70 | n.s. | n.s. | n.s. | n.s. | n.s. | n.s. | n.s. | n.s. |
| | Kolf2.1J | | | | | | | | |
| | A85V | n.s. | n.s. | ≤0.05 | n.s. | n.s. | n.s. | n.s. | n.s. |
| | N141I | n.s. | n.s. | ≤0.001 | ≤0.01 | n.s. | n.s. | n.s. | ≤0.01 |
| | M239I | n.s. | n.s. | ≤0.01 | ≤0.05 | n.s. | ≤0.01 | ≤0.05 | ≤0.001 |
| | **Day 12** | | | | | | | | |
| | ISO AG70 | | | | | | | | |
| | AG70 | n.s. | n.s. | ≤0.01 | n.s. | n.s. | n.s. | n.s. | n.s. |
| | Kolf2.1J | | | | | | | | |
| | A85V | n.s. | n.s. | n.s. | n.s. | n.s. | ≤0.001 | n.s. | ≤0.05 |
| | N141I | ≤0.001 | ≤0.001 | ≤0.05 | ≤0.05 | n.s. | ≤0.05 | n.s. | n.s. |
| | M239I | ≤0.01 | ≤0.01 | n.s. | n.s. | n.s. | ≤0.01 | n.s. | ≤0.01 |
| Amyloid Precursor Protein | **Day 8** | | | | | | | | |
| | ISO KP9 | | | | | | | | |
| | KP9 | n.s. | n.s. | n.s. | n.s. | n.s. | n.s. | n.s. | n.s. |
| | Kolf2.1J | | | | | | | | |
| | E693Q | ≤0.05 | n.s. | n.s. | n.s. | n.s. | n.s. | n.s. | n.s. |
| | V717I | ≤0.001 | ≤0.01 | ≤0.01 | ≤0.05 | n.s. | n.s. | n.s. | ≤0.01 |
| | **Day 12** | | | | | | | | |
| | ISO KP9 | | | | | | | | |
| | KP9 | n.s. | n.s. | n.s. | n.s. | n.s. | n.s. | n.s. | n.s. |
| | Kolf2.1J | | | | | | | | |
| | E693Q | n.s. | n.s. | n.s. | n.s. | ≤0.01 | n.s. | n.s. | n.s. |
| | V717I | ≤0.05 | ≤0.05 | ≤0.05 | n.s. | ≤0.01 | ≤0.05 | n.s. | n.s. |

**Supplemental Table 1. Mitochondrial Stress Test and Electron Flow Assay p-values for PS1, PS2, and APP neurons at DIV8 and DIV12.** Probability values for the Mitochondrial Stress Test (MST) and Electron Flow Assays (EFA) performed using iPSC-derived neurons at DIV 8 and DIV12 shown in Supplemental Figure 3. For both tests, the data were analyzed by comparing the average of 12 technical replicates across 3 independent experiments. Statistical significance was calculated using a t-test for each AD line vs its ISO control. A 2-way ANOVA with Fisher's *post hoc* test was used for Kolf2.1J vs engineered alleles. n.s.=not significant.

| Gene | Mutation | Clone ID | Exon | cDNA | Sequenced Genotype | Passed Testing |
|---|---|---|---|---|---|---|
| PS1 | A79V | A02 | 4 | c.236 C>T | CT | not tested |
| PS1 | A79V | B05 | 4 | c.236 C>T | CT | not tested |
| **PS1** | **A79V** | **B06** | **4** | **c.236 C>T** | **CT** | **Y** |
| PS1 | A79V | B07 | 4 | c.236 C>T | CT | not tested |
| **PS1** | **M139V** | **HET1** | **7** | **c.415 A>G** | **AG** | **Y** |
| PS1 | M139V | HET2 | 7 | c.415 A>G | AG | not tested |
| PS1 | M139V | HET3 | 7 | c.415 A>G | AG | not tested |
| PS1 | M139V | HET4 | 7 | c.415 A>G | AG | not tested |
| **PS1** | **E280A** | **G01.2** | **8** | **c.838 A>C** | **AC** | **Y** |
| PS1 | E280A | B10 | 8 | c.838 A>C | AC | not tested |
| PS1 | E280A | D02 | 8 | c.838 A>C | AC | not tested |
| PS1 | E280A | A11.2 | 8 | c.838 A>C | AC | not tested |
| PS2 | A85V | HET1 | 5 | c.254 C>T | CT | Y |
| PS2 | A85V | HET2 | 5 | c.254 C>T | CT | N |
| PS2 | A85V | HET3 | 5 | c.254 C>T | CT | N |
| **PS2** | **A85V** | **HET4** | **5** | **c.254 C>T** | **CT** | **Y** |
| **PS2** | **N141I** | **A10** | **6** | **c.422 A>T** | **AT** | **Y** |
| PS2 | N141I | C02 | 6 | c.422 A>T | AT | Y |
| PS2 | N141I | D11 | 6 | c.422 A>T | TT | not tested |
| PS2 | N141I | E02 | 6 | c.422 A>T | TT | Y |
| PS2 | M239I | C08 | 8 | c.717 G>A | GA | Y |
| **PS2** | **M239I** | **F04** | **8** | **c.717 G>A** | **GA** | **Y** |
| PS2 | M239I | F06 | 8 | c.717 G>A | GA | Y |
| PS2 | M239I | E05 | 8 | c.717 G>A | GA | not tested |
| APP | E693Q | HET1 | 17 | c.2077 G>C | GC | not tested |
| **APP** | **E693Q** | **HET2** | **17** | **c.2077 G>C** | **GC** | **Y** |
| APP | E693Q | HET3 | 17 | c.2077 G>C | 36bp homozygous del | not tested |
| **APP** | **V717I** | **HET1** | **17** | **c.2148 G>A** | **GA** | **Y** |
| APP | V717I | HET2 | 17 | c.2148 G>A | GA | not tested |
| APP | V717I | HET3 | 17 | c.2148 G>A | GA | not tested |
| APP | V717I | HET4 | 17 | c.2148 G>A | GA | not tested |

**Supplemental Table 2. Kolf2.1J engineered mutant AD clones selected for study.** All early access heterozygous clones received from the iNDI project for potential use are listed. These clones were thawed, expanded, and frozen. Cell pellets from the clones were submitted for pluripotency and karyotype analysis. Passed testing = the clone showed appropriate pluripotency markers and did not have any large chromosomal abnormalities. Genomic DNA was extracted from cell pellets and each mutation was verified via PCR amplification followed by Sanger sequencing. Clones with bolded text were selected for further study. PS1, PS2 and APP codon numbering is based on the NM_000021.4, NM_000447.3, and NM_000484.4 transcript sequences respectively.

| iPSC name | Gene | Mutation | Media | Substrate |
|---|---|---|---|---|
| AG67 | PS1 | A246E | Coriell media* | gelatin and MEF** |
| ISO-AG67 | - | - | E8 Flex | vitronectin |
| CW10192 | - | - | E8 | vitronectin |
| A79V | PS1 | A79V | StemFlex | matrigel |
| M139V | PS1 | M139V | StemFlex | matrigel |
| E280A | PS1 | E280A | StemFlex | matrigel |
| Kolf2.1J | - | - | StemFlex | synthemax II-SC |
| AG70 | PS2 | N141I | Coriell media* | gelatin and MEF** |
| ISO-AG70 | - | - | E8 Flex | vitronectin |
| CW50062 | - | - | E8 | vitronectin |
| A85V | PS2 | A85V | StemFlex | matrigel |
| N141I | PS2 | N141I | StemFlex | matrigel |
| M239I | PS2 | M239I | StemFlex | matrigel |
| KP9 | APP | K670N/M671L | StemMACs iPS-Brew XF | matrigel |
| ISO-KP9 | - | - | E8 Flex | vitronectin |
| E693Q | APP | E693Q | StemFlex | matrigel |
| V717I | APP | V717I | StemFlex | matrigel |

**Supplemental Table 3. Original culture conditions of iPSC used in this study.** *Coriell media has a base of DMEM/F12 and final concentrations of the following components: 20% knock-out serum replacement, 2mM L-glutamine, 1x non-essential amino acids, 0.1mM 2-mercaptoethanol, 10ng/mL bFGF (fibroblast growth factor – basic). **MEF = mouse embryonic fibroblasts.
